# Back to the holobiont: ecophysiological and systemic responses of rooted-cuttings inoculated with a synthetic community

**DOI:** 10.1101/2023.11.02.565299

**Authors:** Marco Sandrini, Walter Chitarra, Chiara Pagliarani, Loredana Moffa, Maurizio Petrozziello, Paola Colla, Raffaella Balestrini, Luca Nerva

## Abstract

Despite microbe-based products for grapevine protection and growth improvement are already available, a few of them contain microbes directly isolated from vine tissues. For this reason, a collection of endophytic bacterial isolates obtained directly from grapevine woody tissues has been used for producing an *ad-hoc* inoculum. The selected bacterial isolates were tested in biocontrol assays against some of the main grapevine pathogens (*e.g.*, *Botrytis cinerea*, *Guignardia bidwellii*, *Neofusicoccum parvum*) and the best performing ones were screened for plant growth promoting (PGP)-traits (*e.g.*, phosphorous solubilization, indole-acetic acid and siderophore production). Before being planted, rooted cuttings were inoculated with two different synthetic communities: the first one was an *ad-hoc* developed microbial community (SynCom), whereas the second was a commercial consortium formed by arbuscular mycorrhizal fungi (AMF) and a rhizosphere bacterial strain (AMF+B). Physiological parameters were monitored to evaluate effects on plant performances, and samples for biochemical and molecular analyses were collected. Integration of physiological data with metabolite profiling and transcriptome sequencing highlighted that the SynCom treatment shaped the plant growth-defence trade-off, by regulating photosynthesis and diverting energy sources towards the activation of defence metabolic pathways. On the other hand, the AMF+B treatment led to a more balanced growth-defence trade-off, though a mild activation of defence mechanisms was also observed in these plants. Our findings suggest that an experimental approach considering both the features of associated microbes and their impacts on plant growth and defence could shed light on the “dark-side effects” of SynCom application, thereby enabling their exploitation with a refined awareness.

## Introduction

Plants naturally share their environment with a multitude of microbes, some of which can colonize their inner tissues becoming endophytes and strongly influencing plants life cycle and responses to the environmental stimuli (Nerva et al., 2022a). The recruitment of microbes by plants (*i.e.*, microbiome assembly) strictly depends on the interaction between plants and the surrounding environment and, once the relationship is established, the plant and its microbiome behave as a unique super organism, referred to as holobiont (Vandenkoornhuyse et al., 2015).

Based on the global climate changing effects and the resulting prediction of more frequent biotic and abiotic stressful events, the development of novel, sustainable crop protection strategies is extremely urgent to improve agricultural resilience (Chitarra et al., 2015; Giudice et al., 2021; Giudice et al., 2022). In this context, a better understanding of plant-microbiome interactions could help to dissect key factors involved in the recruitment of beneficial microbes by the host (Nerva et al., 2022b). Thanks to the great advances in microbial biotechnology, metagenomics and the extensive collections of microbes recently developed, we have a treasure at our disposal to manipulate bacterial communities on a large scale (Zou et al., 2019; Sandrini et al., 2022). In this fashion, it is possible to cultivate pure characterized strains and develop synthetic microbial communities (SynComs) to mimic natural microbiome functions and study, through multi- and interdisciplinary approaches, the plant microbiome assembly and its effects on plant performances under diverse environmental conditions (Sandrini et al., 2022). Interestingly, the assembly rules for establishing plant microbiota have been revealed in gnotobiotic *Arabidopsis* plants using a drop-out and a late introduction approach with SynComs composed by 62 native strains. The authors found that community assembly has historical contingency with priority effects and a certain degree of resistance to late microbe arrivals (Carlström et al., 2019). These findings provided new insights into plant-microbiome interactions and suggested that, to be successful, application of SynComs should be performed at the early stages of the host life cycle, when the microbiome is still under development.

As cited before, many beneficial endophytes-inhabiting plant tissues (*e.g.*, Plant Growth-Promoting microorganisms, as for instance *Actinomycetes*, and Arbuscular Mycorrhizal Fungi–AMF) play key roles in protecting plants against biotic and abiotic stresses (Chitarra et al., 2016; Carrión et al., 2019). Besides basic research to study the complex host-microbiome interactions, SynComs can be exploited to promote growth and other beneficial traits in the host. During the last decade, many SynCom approaches have been reported in literature (most of them conducted in axenic conditions), highlighting the increased attention to this subject for orienting future sustainable agricultural practices (Sandrini et al., 2022).

Plant diseases cause significant losses in agricultural production that lead not only to lower yields and decreased quality, but also to loss of biodiversity, mitigation costs due to control measures, and an important downstream impact on human health (Ristaino et al., 2021). Plants possess natural defence mechanisms that are finely regulated by several phytochemicals. Once infected by pathogens, plants promptly react activating innate defence mechanisms that are tightly mediated by jasmonic acid (JA), ethylene (Et) and salicylic acid (SA). These phytohormones orchestrate several signalling pathways from cells to systemic routes by means of the so-called systemic acquired resistance (SAR, mediated by SA) or induced systemic resistance (ISR, mediated by JA and Et) (Burketova et al., 2015). The latter can be activated in consequence to pathogen attacks but also by soil-inhabiting beneficial microbes (*e.g.*, PGPB, AMF) recruited by plants thanks to the modulation of signalling root exudates abundantly released under stressful conditions (Berendsen et al., 2018). Thus, the development of tailored SynComs could represent a powerful tool to prime plants against biotic (and/or abiotic) stress, preventing food losses. Indeed, Berendsen et al. (2018) developed a simplified SynCom formed by three bacterial species that synergistically protected *Arabidopsis* plants against downy mildew. In another study, by using a SynCom composed by 38 bacterial strains, Lebeis (2015) demonstrated that immune signalling is the driver of microbiome development in *Arabidopsis*. Recently, Li et al. (2021) assembled two SynComs (one complex and another one simplified to 4-species forming community) with disease controlling functions against *Fusarium* sp., the causal agent of root rot disease in *Astragalus mongholicus*. These authors observed that both SynComs were successful in controlling the disease development *via* synergistic cooperation by activating ISR paths in the host.

It is worth noting that the native microbiome is continuously modulated based on the host genotype and on environmental stimuli. Similarly, SynCom structure and functionality can be strongly influenced by the same variables, although further studies are needed. A better understanding of SynCom functionality in “natural” environments would allow to fully exploit their potential (Wei et al., 2018; Veach et al., 2019), especially considering that, many of the studies available to date were conducted in axenic conditions.

Grapevine represents one of the most important fruit crops worldwide, hardly infected by many pathogens (mainly fungal ones) both in pre- and post-harvest (*e.g.*, powdery mildews, grey mold, esca syndrome). These pathogens cause important damages, usually controlled by massive pesticide application that strongly impact agroecosystems, beneficial microbiota and human health, making urgent the development of new sustainable alternatives (Armijo et al., 2016; Nerva et al., 2019; Giudice et al., 2022; Nerva et al., 2022a). Therefore, in this study we aimed at investigating whether the plant inoculation with a simplified SynCom formed by biocontrol agents could trigger systemic defence responses in the host. To address the above question, a SynCom formed by seven bacterial isolates retrieved from the inner grapevine woody tissues (Nerva et al., 2022a) was developed and inoculated in grapevine rooted cuttings prior to planting them in field. The newly developed SynCom was compared with a commercial one formed by a mixed inoculum of AMF and a bacterial strain. Afterwards, combined ecophysiological, biochemical and molecular approaches were used to investigate the SynCom effects on the host physiological performances, plant growth promotion and ISR activation. Criticisms and suggestions for the implementation of SynCom protocols for future scale up and field applications have also been discussed.

## Results

### Isolation and molecular characterization of the bacterial collection

Forty-three bacterial isolates were collected, and the molecular identification was achieved by 16S rRNA gene sequencing analysis. Forty-four isolates were identified, most of which belonging to the Actinobacteria phylum (Table 1). In fact, although a specific protocol for Actinobacteria isolation was adopted, 22 out of the 44 isolates were found to be Actinobacteria, and 22 out of 44 were Proteobacteria (Table 1). The 16S sequences of each isolate were deposited in NCBI GenBank under the accessions OP994307-OP994344. Checking in literature, no one among the bacterial isolates showed similarities with those harmful for humans or plants.

**Table 1.**
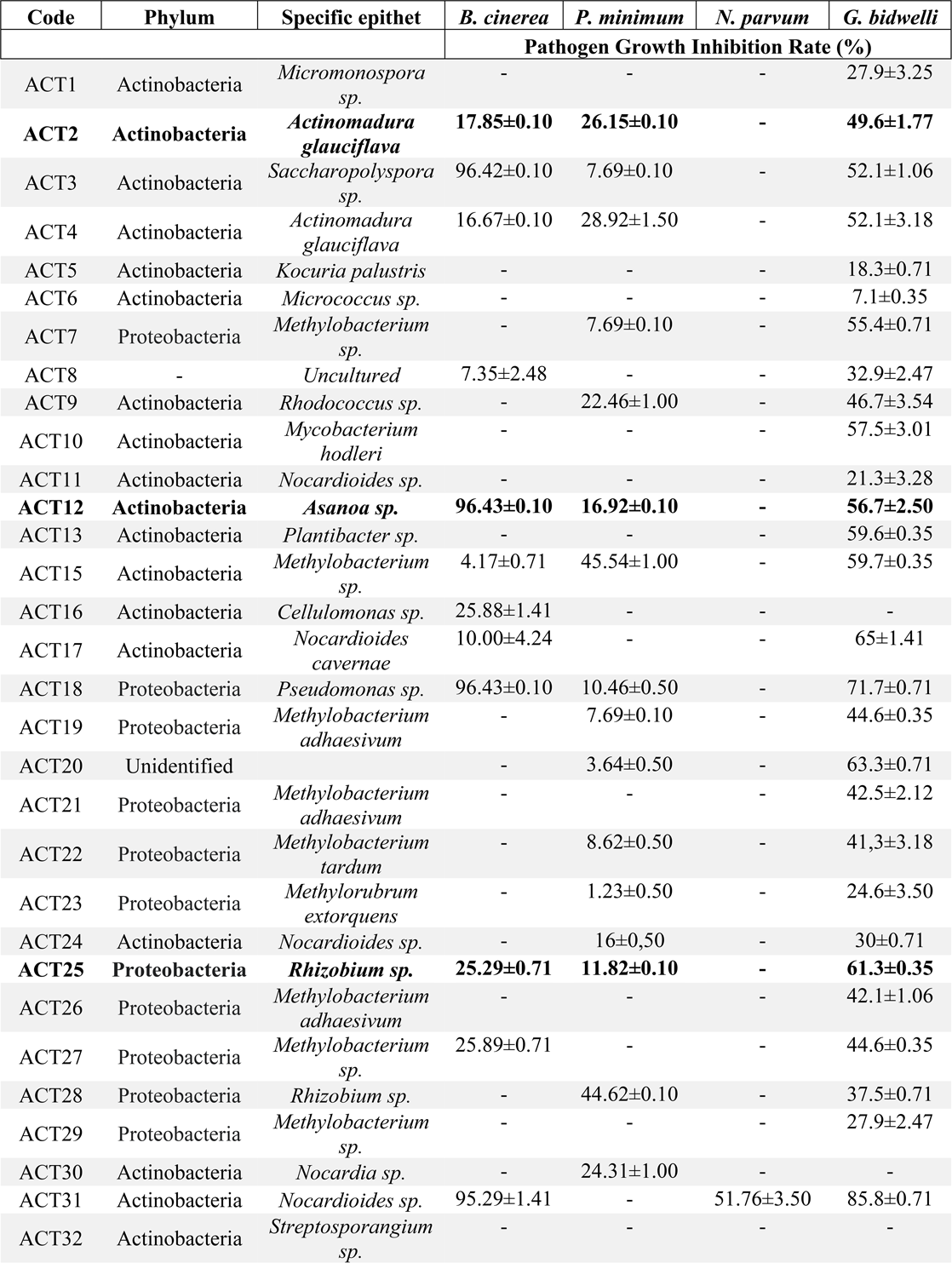
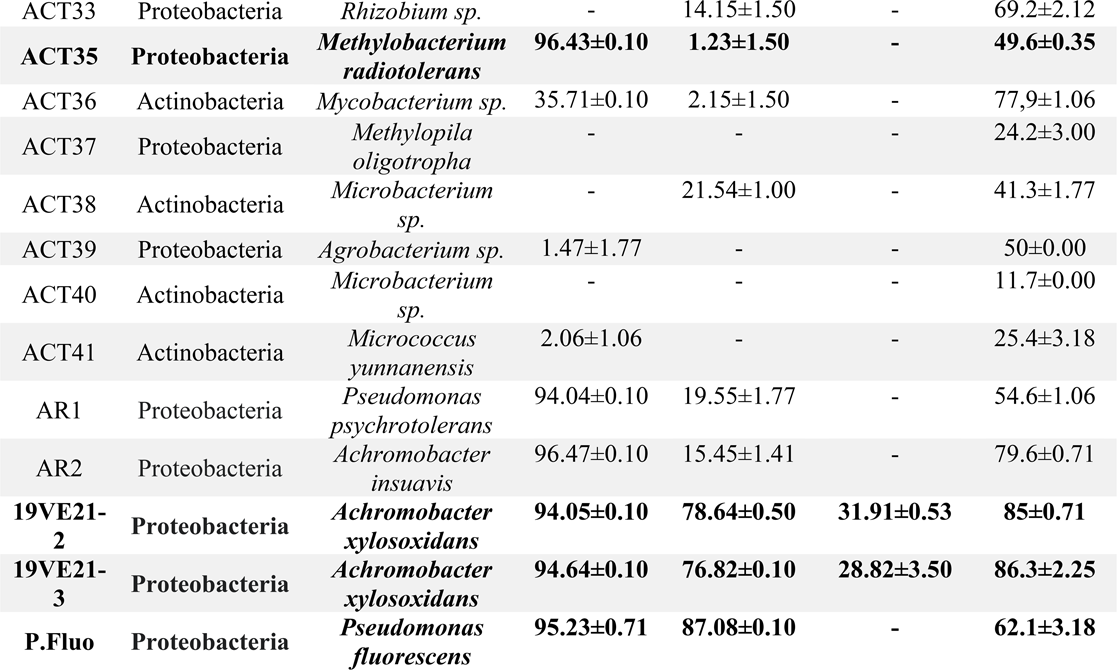
Biological control activity of the whole bacterial collection. Data for biological control activity are referred to the pathogen growth inhibition rate (%) toward each fungal pathogen. The reported values of pathogen growth inhibition rate are the mean values of three replicates ± SD for each isolate. The selected isolates forming the SynCom are reported in bold.

Since the purpose of this survey went beyond the establishment of a collection of grapevine endophytes, the whole 44 bacterial isolates were *in vitro* assessed for biocontrol activity against some of the most important and widespread grapevine pathogens (Table 1) and for PGP-traits (Supplementary Table S1), as below reported.

### *In vitro* evaluation of biocontrol and Plant Growth Promoting (PGP) activities of the bacterial isolates

In the biocontrol Petri dish assay, 44 strains showed a different degree of pathogen containment and the best performing isolates, considering all pathogens and the ability to grow together, were selected as good candidates to build a SynCom (Table 1 and Figure 1). Particularly, an antagonistic activity towards at least three pathogens and a pathogen growth inhibition rate higher than 45%, facing at least one fungus, was adopted as selection criteria. The seven selected bacterial strains are highlighted in bold in Table 1. Since the high efficacy in controlling pathogens and the potential production of volatile organic compounds (VOCs), the seven isolates were challenged in septate Petri dishes, though none of them showed the ability to inhibit the pathogen in such experimental set-up (data not shown). Further analyses were performed to evaluate the ability of SynCom candidate members to stimulate plant growth and abiotic stress tolerance (Supplementary Table S1): all the seven candidates were able to solubilize starch, with the 19VE21-2 isolate showing the largest halo zone around the colonies. Four isolates out of the seven were able to solubilize phosphate, with 19VE21-2 showing the wider clear zone around the colonies. Three isolates were able to grow on NFb agar plate thus showing a potential to fix nitrogen, and among those ACT2 showed the most preeminent growth. Five isolates were proved to be siderophore producers, with ACT25 and 19VE21-3 showing the widest yellow halo appearance around bacterial colonies on the Chrome Azurol medium. Five isolates were identified as IAA producers, and P. Fluo was the one showing the most abundant IAA production. Four strains were also able to degrade ACC, and particularly the ACT35 and 19VE21-2 isolates displayed the highest consumption activities. As the last PGP-related trait, we evaluated the ability of the selected isolates to grow in presence of NaCl at different concentrations (0, 1.5 and 3% w/v), and we found that six out of the seven candidates displayed an enhanced salinity tolerance. The biocontrol activity and the PGP-traits are reported in Fig. 1 for each of the selected isolates along with pictures taken at the end of the biocontrol assay on *Guignardia bidwellii* (the causal agent of black rot).

**Figure 1.**
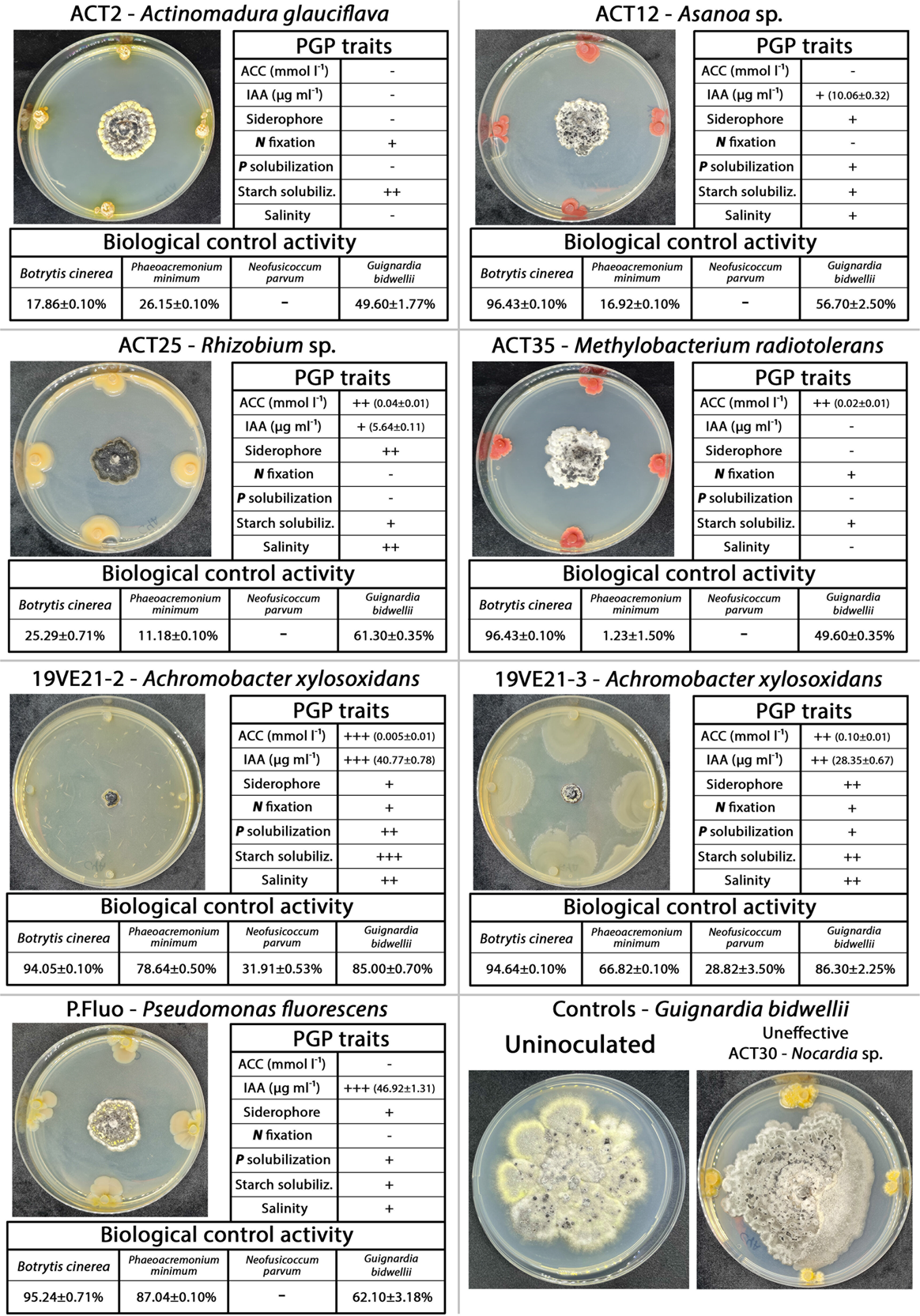
Plant growth promotion and biological control traits of the selected isolates forming the SynCom. For PGP traits the symbol + represents the presence of growth/halo, whereas the symbol – represents negative results for the test. Data for 1-aminocyclopropane-1-carboxylate (ACC) deaminase activity: +++, good; ++, medium; +, slight; values are the mean ± SD. Data for IAA production: +, quantity < 20 µg ml^-1^; ++, quantity between 20 and 40 µg ml^-1^; +++, quantity >40 µg ml^-1^; values are the mean ± SD. Data for siderophore production: +, zone of yellow halo up to 2 mm; ++, zone of yellow halo bigger than 2 mm up to 5 mm. Data for *N* fixation: +, growth capability on Nfb agar; -, absence of growth. Data of *P* solubilization: +, isolates with halo diameter up to 1 mm; ++, halo diameter bigger than 1 mm up to 5 mm. Data for starch hydrolysis: + (0 cm up to 0.4 cm), ++ (0.4 cm up to 1 cm), +++ (higher than 1 cm) mean low, medium and high halo diameter on the medium after incubation, respectively. Data related to salinity tolerance: +, low growth; ++, medium growth; +++, good growth. For each PGP trait, at least three replicates per isolate were considered. Data for biological control activity are referred to the % of growth inhibition rate toward each selected fungal pathogen. Insets: examples of biocontrol screening test conducted *in vitro* for each selected isolate forming the SynCom.

Finally, to limit reciprocal inhibition effects, a strain compatibility assay was performed. The results revealed a good aptitude of the candidates to live together, thereby confirming them as promising members to form a grape-specific synthetic community (Fig. S1).

### Analysis of AM root colonization

To confirm the establishment of mycorrhizal symbiosis on roots of rooted cuttings inoculated with the commercial mixed formulation containing both AMF species and a bacterial strain (AMF+B), the percentage of arbuscules in root cortical cells was calculated on three randomly selected plants for each treatment. Microscopic observations of stained roots revealed the presence of mycorrhizal structures at different extents based on the treatment. The roots of AMF+B treated plants had a significantly higher percentage of mycorrhization frequency (F), ranging around 100%; intensity of the mycorrhizal colonisation in either the root system (M) or fragments (m) ranged around 60%; the arbuscule abundance ranged around 94% in the root fragments (a) and around 54% in the root system (A) (Fig. 2a). Conversely, very few fungal structures were observed in the roots collected from SynCom and CTRL samples (Fig. 2a). Additionally, to check the functionality of AM symbiosis, the expression analysis of three AM symbiosis-related genes (*i.e.*, *VvCCD7* and *VvCCD8*, both involved in strigolactone biosynthesis, and *VvPT1-3*, encoding a grape phosphate transporter) was performed. While *VvCCD7* was significantly up-regulated in AMF+B- and SynCom-treated plants with respect to CTRL plants (Fig. 2b), transcription of *VvCCD8* did not significantly vary among treatments (Fig. 2c). The *VvPT1-3* gene was previously reported in grapevine as a marker of functional AM symbiosis establishment (Balestrini et al., 2017; Nerva et al., 2021a). Here, the expression profile of this gene was significantly modulated by AMF+B, showing a strong up-regulation upon this treatment with respect to both CTRL and SynCom samples (Fig. 2d).

**Figure 2.**
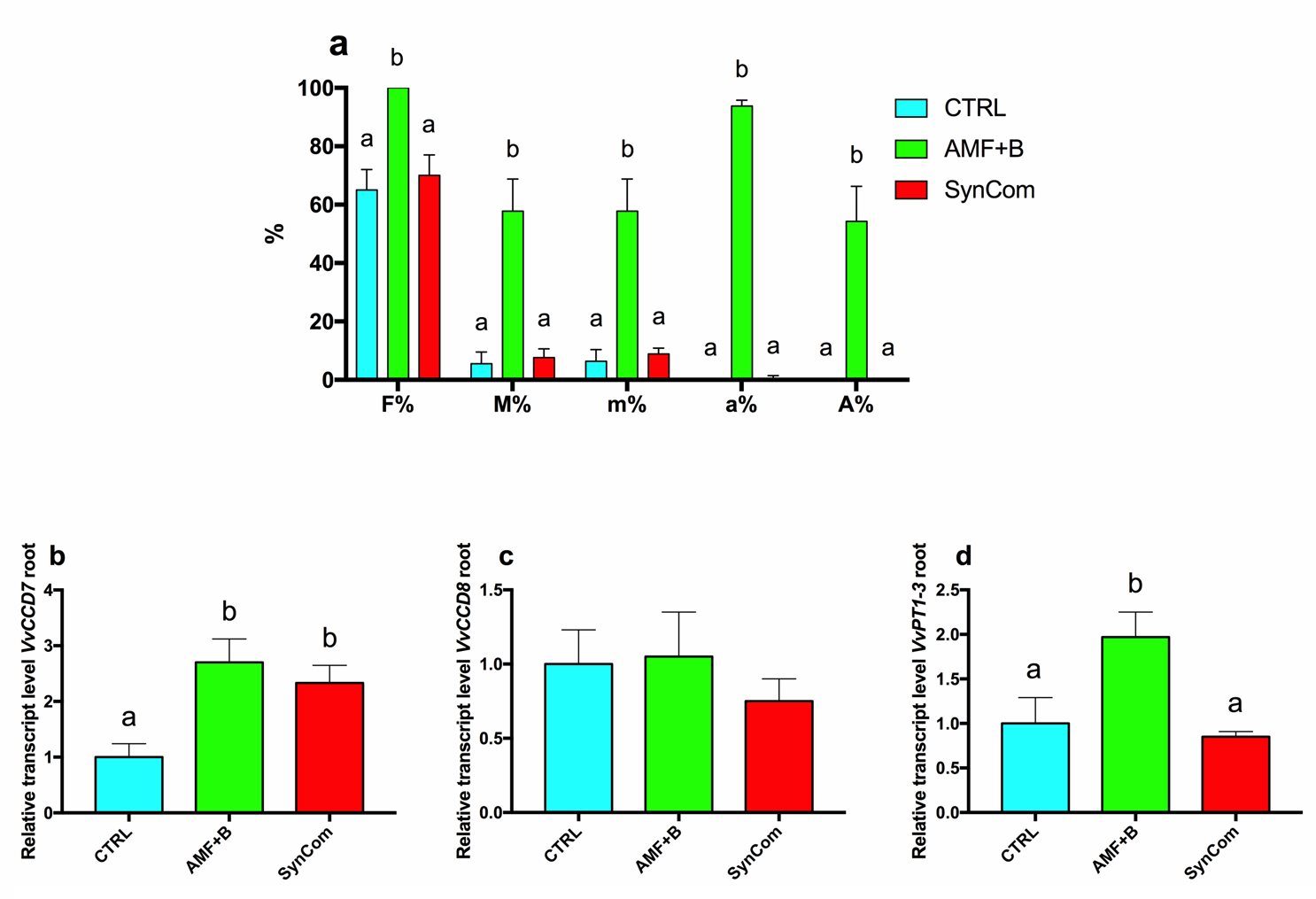
AMF colonization in grapevine roots. **a)** Colonization rate in CTRL, AMF+B and SynCom treatments. Data are expressed as mean ± SD (n = 3). F%, Frequency of mycorrhization in the root system; M%, Intensity of the mycorrhizal colonization in the root system; m%, Intensity of the mycorrhizal colonization in the root fragments; a%, Arbuscule abundance in mycorrhizal parts of root fragments; A%, Arbuscule abundance in the root system. **b-d**) Expression profiles of the strigolactone biosynthesis-related genes *VvCCD7* and *VvCCD8*, respectively (**b, c)**, and of a grapevine phosphate transporter-encoding gene (**d**), *VvPT1-3*, marker of functional symbiosis. Lowercase letters above bars denote significant differences attested by Tukey’s HSD test (*P* ≤ *0.05*). CTRL, control plants; AMF+B, commercial AMF + Bacteria mixed inoculum-treated plants; SynCom, Synthetic Community-treated plants.

Collectively, these results suggested that AMF+B roots were efficiently colonized, and that AM symbiosis was successfully established in the grapevine cuttings. Conversely, colonization from native AMF in SynCom and CTRL plants was not relevant.

### Microbiome analysis of root-associated endophytes

Bacterial and fungal community were analysed at the genus level: after chimera removal and taxonomical assignment, the number of retained sequences was always above 30.000 (detailed results of sequencing are reported in Supplementary Table S3). Shannon index diversity indicated that AMF+B-treated samples showed a significantly higher bacterial diversity in comparison to both CTRL and SynCom, while SynCom did not differ from CTRL (Supplementary Table S4). Shannon index for the fungal community attested that all treatments significantly differed from each other. Specifically, samples subjected to the AMF+B treatment exhibited the highest diversity, followed by CTRL, and SynCom (Supplementary Table S4). The non-metric multidimensional scaling (NMDS), based on Bray-Curtis dissimilarity matrixes, highlighted that both bacterial (Fig. S2a) and fungal communities (Fig. S2b) were significantly influenced by the applied treatment. These results were also confirmed by PERMANOVA analysis, from which a significant distinction emerged between bacterial and fungal communities in all tested conditions (Fig. S2).

The composition of the bacteria community at the genus level is reported for each sample type in Supplementary Table S5. A graphic distribution of the taxa representing at least the 1% of the community is displayed in Fig. 3a. The comparison of the bacterial community among treatments revealed that the genus *Pelosinus* (phylum Firmicutes) was negatively affected by the inoculation of both AMF+B and SynCom. On the contrary, the abundance of the genus *Flavobacterium* (phylum Gammaproteobacteria) was positively modulated by both AMF+B and SynCom treatments. In parallel, amplicon sequence variant (ASV) belonging to the genera inoculated with the SynCom (*i.e.*, *Pseudomonas*, *Asanoa*, *Achromobacter*, *Actinomadura*, *Rhizobium* and *Methylobacteria*) were significantly overrepresented in the SynCom samples (Supplementary Table S5).

**Figure 3.**
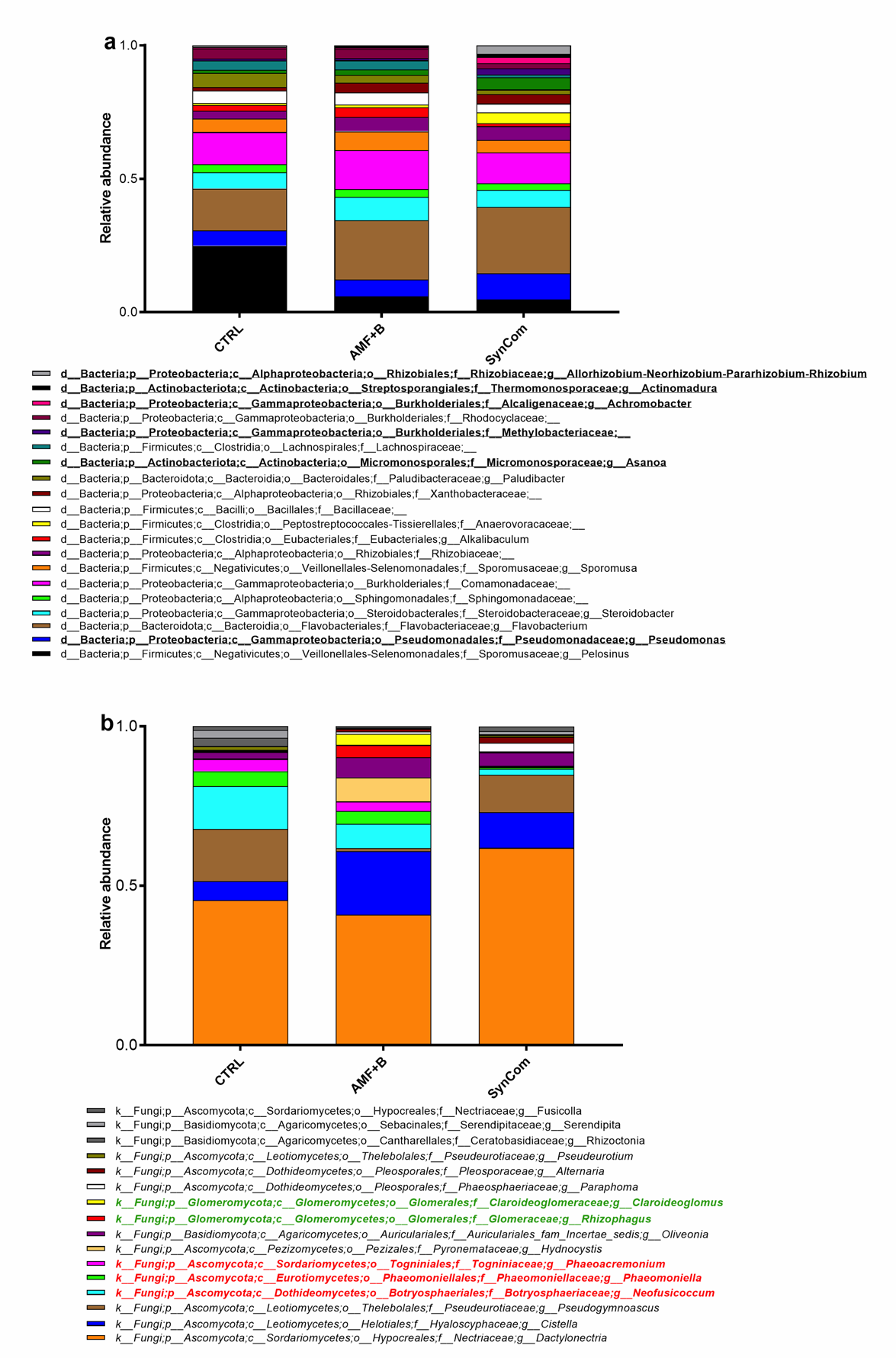
Relative abundances of bacterial and fungal genera. a) Bacterial genera added with SynCom inoculation are highlighted in bold. Only genera representing at least the 1% over the total of classified amplicons were retained. b) Fungal genera belonging to mycorrhizal species added with the commercial formulation (AMF+B) are highlighted in green. Wood decay-associated fungal genera are highlighted in red. Only genera representing at least the 1% over the total of classified amplicons were retained.

The fungal community composition at the genus level is reported in Supplementary Table S6 for each sample type. A graphic outline of the taxa representing at least the 1% of the community is shown in Fig. 3b. A further comparison of the fungal community among treatments revealed interesting outputs. The *Claroideoglomus* and *Funnelliformis* genera, both belonging to the Glomeromycota phylum, were overrepresented only in the AMF+B samples where mycorrhizal fungi were added. In parallel, the *Neofusicoccum*, *Phaeomoniella* and *Phaeoacremonium* genera, commonly associated with wood decay, were underrepresented in the SynCom-treated plants with respect to AMF+B and CTRL plants (Supplementary Table S6).

The network analysis of microbial communities, performed among the different treatments and conducted using the co-occurrence method, is detailed in Supplementary Table S7 and depicted in Fig. S3. Co-occurrence networks were calculated by combining data from 16S and ITS sequencing analyses for CTRL (Fig. S3a), AMF+B (Fig. S3b) and SynCom (Fig. S3c). Additionally, root biochemical measurements were incorporated as features during the analysis process. Interestingly, when the whole dataset was analysed, the co-occurrence network (Fig. 4) highlighted a significant mutual exclusion effect between some of the bacteria inoculated with the SynCom and wood fungal pathogens (Supplementary Table S7 – sheet d and Fig. 4). Among these interactions, the *Asanoa* genus displayed a mutual exclusion correlation with all the wood fungal pathogens previously considered (*i.e.*, genera *Neofusicoccum*, *Phaeomoniella* and *Phaeoacremonium*). On the contrary, the fungal pathogens were all linked by a copresence correlation.

**Figure 4.**
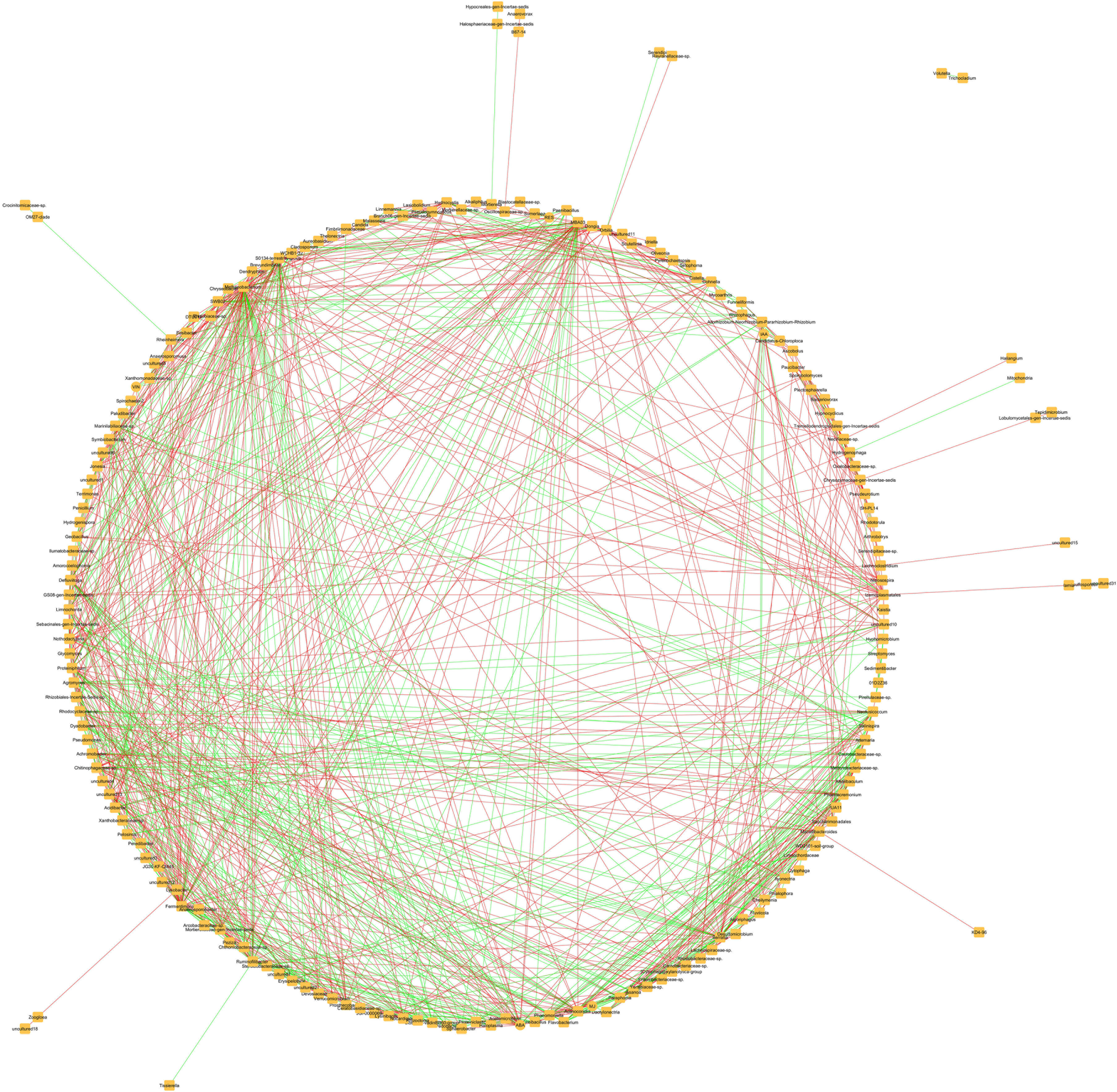
Network co-occurrence analysis for the whole dataset. Network analysis was performed by using as input the microbial abundance data (bacterial and fungi combined) and the biochemical measurements of CTRL, AMF+B and SynCom, and analysing them all together. Green lines point to copresence interactions, while red lines highlight mutual exclusion. Each node represents a taxonomic group.

### Outline of physiological and biochemical responses in field conditions

The leaf gas exchange rates were measured in treated vines to establish whether the inoculation with SynCom was able to alter/improve the plants’ physiological performances in open field conditions in comparison with the other treatment (AMF+B) and with the CTRL. Collected data indicated that the rooted cuttings previously inoculated with AMF+B had a significantly higher Net Photosynthesis (Pn) when compared to CTRL and SynCom-treated plants, whereas the latter showed the significantly lowest Pn values when compared both to AMF+B and CTRL treatments (Fig. 5a). The rates of stomatal conductance (g_s_) significantly increased following the AMF+B treatment with respect to the other tested conditions. A slight, though not significant rise in g_s_ was also observed in SynCom-treated vines in comparison with CTRL (Fig. 5b). Conversely, the values of substomatal CO_2_ concentration (Ci) did not significantly change in the analysed plants, independently of the treatment (Fig. 5c). Additionally, the Apparent Carboxylation Efficiency (ACE, calculated as Pn Ci^-1^ ratio) and intrinsic Water Use Efficiency (iWUE, calculated as Pn g_s_^-1^ ratio) were calculated, highlighting significantly lower values in the SynCom-treated plants with respect to all other conditions (Fig. 5d,e). Overall, the analysis of ecophysiological parameters pointed to a marked photosynthetic imbalance in the SynCom-inoculated vines.

**Figure 5.**
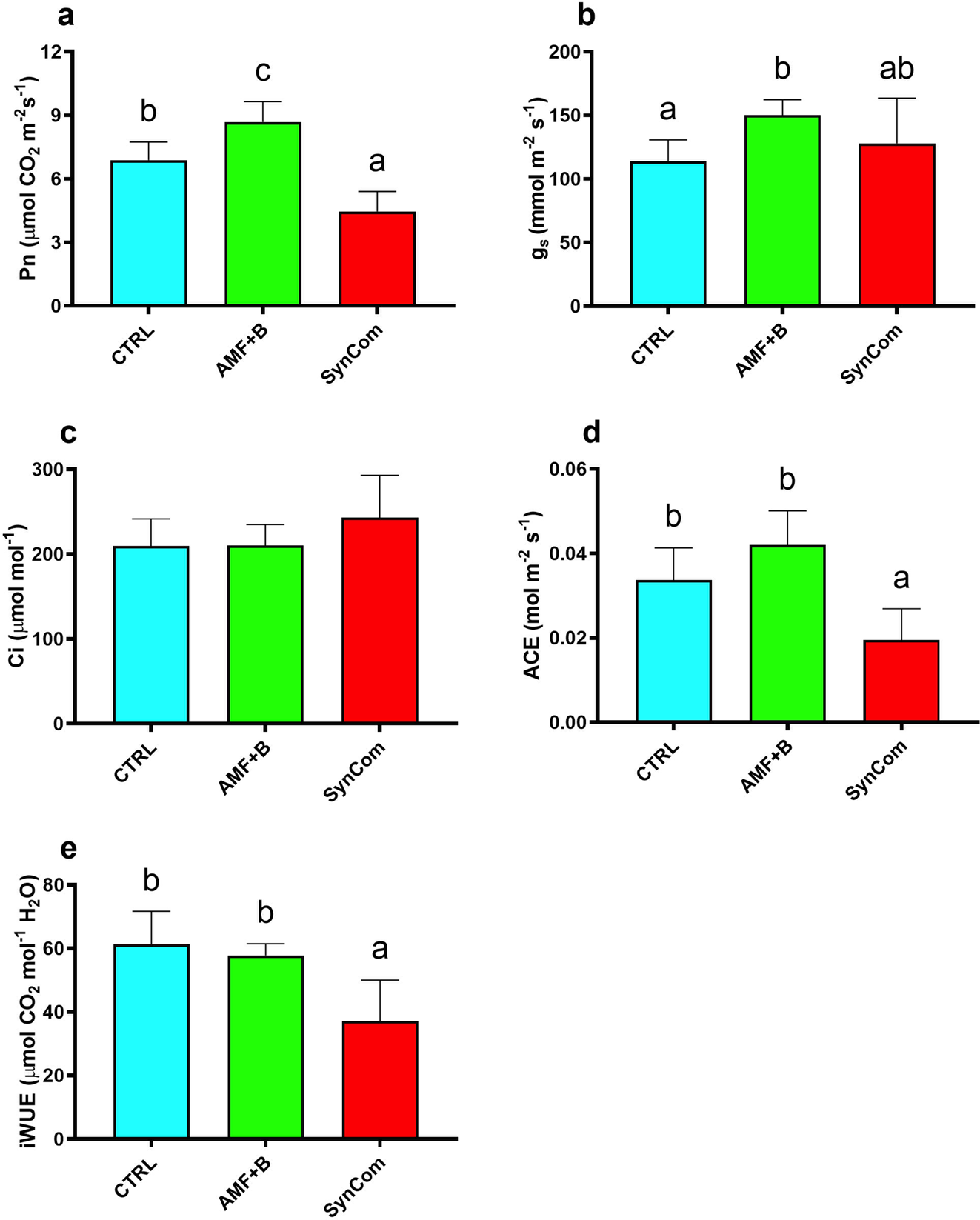
Instantaneous leaf gas exchange measurements. Values of **a)** net photosynthesis (Pn); **b)** stomatal conductance (g_s_); **c)** intercellular CO_2_ concentration (Ci); **d)** apparent carboxylation efficiency (ACE); and **e)** intrinsic water use efficiency (iWUE) recorded in CTRL (control), AMF+B, (commercial AMF + Bacteria mixed inoculum)-treated and SynCom (Synthetic Community)-treated plants. Data are expressed as mean ± SD (n = 36). Different lowercase letters above bars indicate significant differences according to Tukey’s HSD test (*P* < *0.05*).

Changes in endogenous biochemical signals occurring in both leaves and roots were also inspected by analysing the accumulation of phytohormones and secondary metabolites involved in plant growth (*i.e.*, IAA) and defence processes (*e.g.*, ABA, *t*-resveratrol, viniferin, methyl salicylate, methyl jasmonate and jasmonate).

Concentration of abscisic acid (ABA) in the leaf significantly decreased in both inoculated plants (AMF+B and SynCom) with respect to CTRL. However, in the SynCom-treated vines the reduction in ABA levels was more pronounced, with values almost half of those quantified in CTRL leaves (Fig. 6a). The analysis of the hormone content in the root highlighted an opposite condition, with significantly higher concentrations in both AMF+B and SynCom-inoculated plants compared to the CTRL (Fig. 6b). Although indol acetic acid (IAA) amounts did not significantly change in the leaf independently of the treatment (Fig. 6c), in roots they were higher in the SynCom-inoculated plants than in AMF+B-treated vines (Fig. 6d). Accumulation of defensive secondary metabolites was overall more accentuated in both leaves and roots of inoculated plants. In detail, leaf samples collected from the SynCom-treated plants showed the highest *t*-resveratrol concentrations, up to three folds than those measured in CTRL leaves (Fig. 6e). Compared with CTRL samples, *t*-resveratrol production was strongly elicited in the roots of inoculated plants, with similar values between SynCom and AMF+B-treated plants (Fig. 6f). Viniferin amounts were almost undetectable in the leaf of not treated vines, whereas they significantly increased following SynCom and AMF+B inoculation (Fig. 6g). A similar result was obtained from the analysis of the metabolite concentration in root tissues (Fig. 6h).

**Figure 6.**
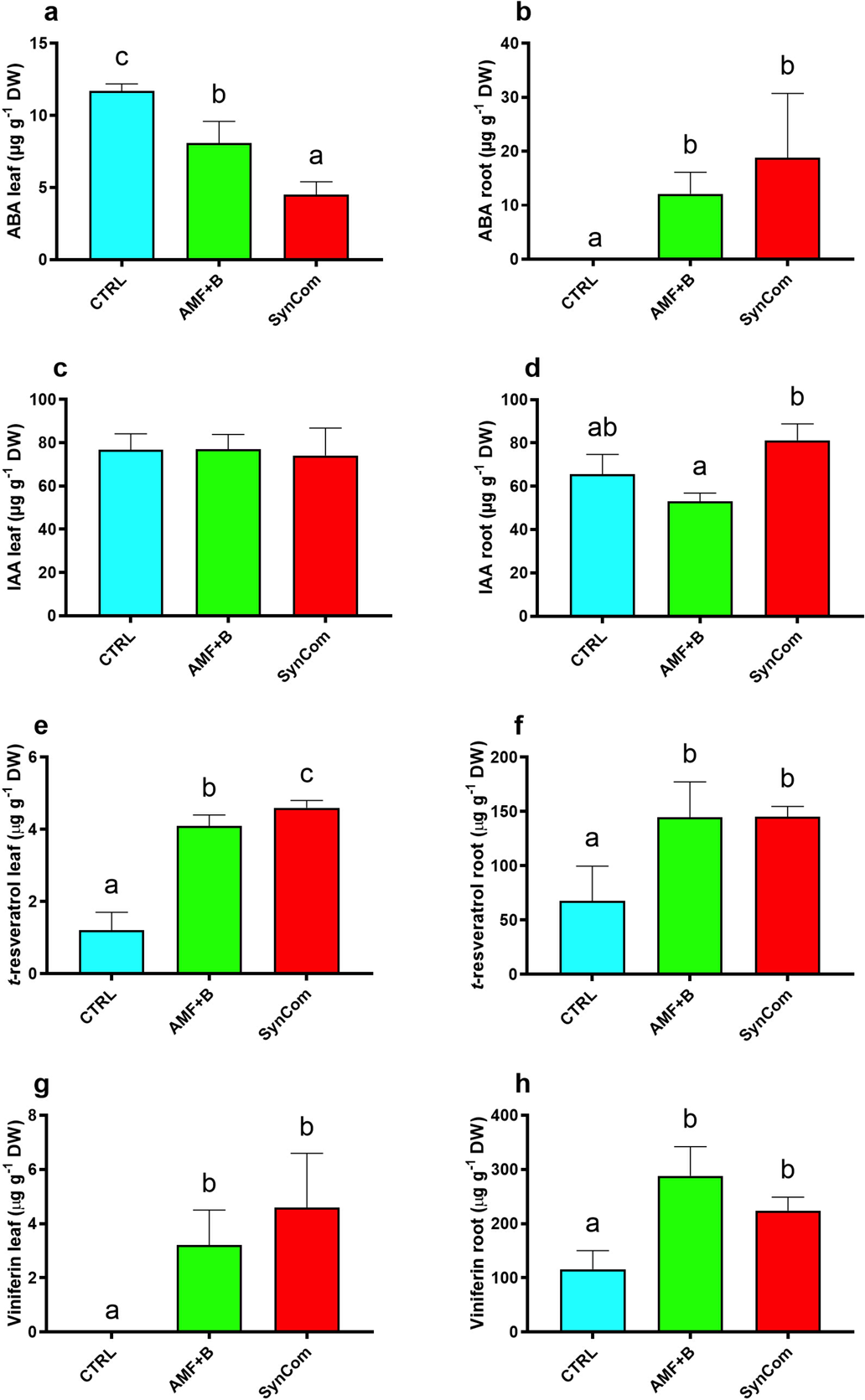
Concentrations of target metabolites in leaf and root tissues. **a,b)** Abscisic acid (ABA) quantification in leaf and root tissues, respectively. **c,d)** Indolacetic acid (IAA) quantification in leaf and root tissues, respectively. **e,f)** *trans*-resveratrol (*t*-resveratrol) quantification in leaf and root tissues, respectively. **g,h)** Viniferin quantification in leaf and root tissues, respectively. Data are expressed as mean ± SD (n = 3). Different lowercase letters above bars indicate significant differences according to Tukey’s HSD test (*P* < *0.05*). CTRL, control plants; AMF+B, commercial AMF + Bacteria mixed inoculum-treated plants; SynCom, Synthetic Community-treated plants.

Additionally, along with the previously mentioned stilbenoids, changes in the content of other core components of the ISR immune system were investigated. To this aim, methyl jasmonate (MJ), and jasmonate (JA) were quantified in the target tissues by using a Gas Chromatography-Mass Spectrometry (GC-MS) approach. Additionally, methyl salicylate (MS), a key component of SAR, was analyzed (Fig. S4). The obtained data indicated that MS and JA levels were not influenced by the adopted treatments. In fact, the concentrations of these compounds did not significantly differ among the tested conditions both in root and leaf tissues (Fig. S4a-b). Although MJ levels in the leaf did not vary among the diverse treatments (Fig. S4a), a significant increase was observed in the root of SynCom-inoculated vines with respect to CTRL plants (Fig. S4b).

### Analysis of candidate gene expression profiles

To complete the characterization of SynCom effects on field-grown vines, changes in the transcript levels of candidate genes involved in defence and growth pathways were investigated in all tested conditions. Starting with the genes linked to establishment of defence mechanisms, the expression of *VvPAL*, encoding a phenylalanine amino lyase, was significantly higher in both leaves and roots of AMF+B- and SynCom-treated plants with respect to CTRL (Fig. 7a,b). A similar expression pattern was observed for *VvSTS1*, encoding a grapevine stilbene synthase, which transcription was strongly induced in the leaves of inoculated plants, regardless of the specific treatment (Fig. 7c). Nonetheless, an opposite result was obtained for the analysis of the same gene in roots. *VvSTS1* expression levels were indeed lower in root samples collected from AMF+B- and SynCom-inoculated vines than CTRL, though the observed downregulation was statistically significant only in AMF+B roots (Fig. 7d). Looking at key players of the plant’s immunity system, both *VvLOX*, encoding a lipoxygenase, and *VvNPR1*, a non-expressor of pathogenesis-related genes 1, are well known genes respectively involved in the onset of ISR and SAR. Independently of the tissue, transcription of *VvLOX* was significantly induced exclusively upon SynCom treatment, showing expression values up to 2-fold higher than AMF+B and CTRL samples (Fig. 7e-f). No significant differences among the treatments were instead observed for *VvNPR1* in leaves (Fig. 7g), whereas a significant up-regulation of this gene occurred in the roots of both AMF+B and SynCom-treated plants with respect to CTRL (Fig. 7h). Compared with AMF+B inoculation, the treatment with SynCom seemed to exert a stronger (though not significant) impact on the modulation of *VvNPR1* expression levels in the vine root (Fig. 7h).

**Figure 7.**
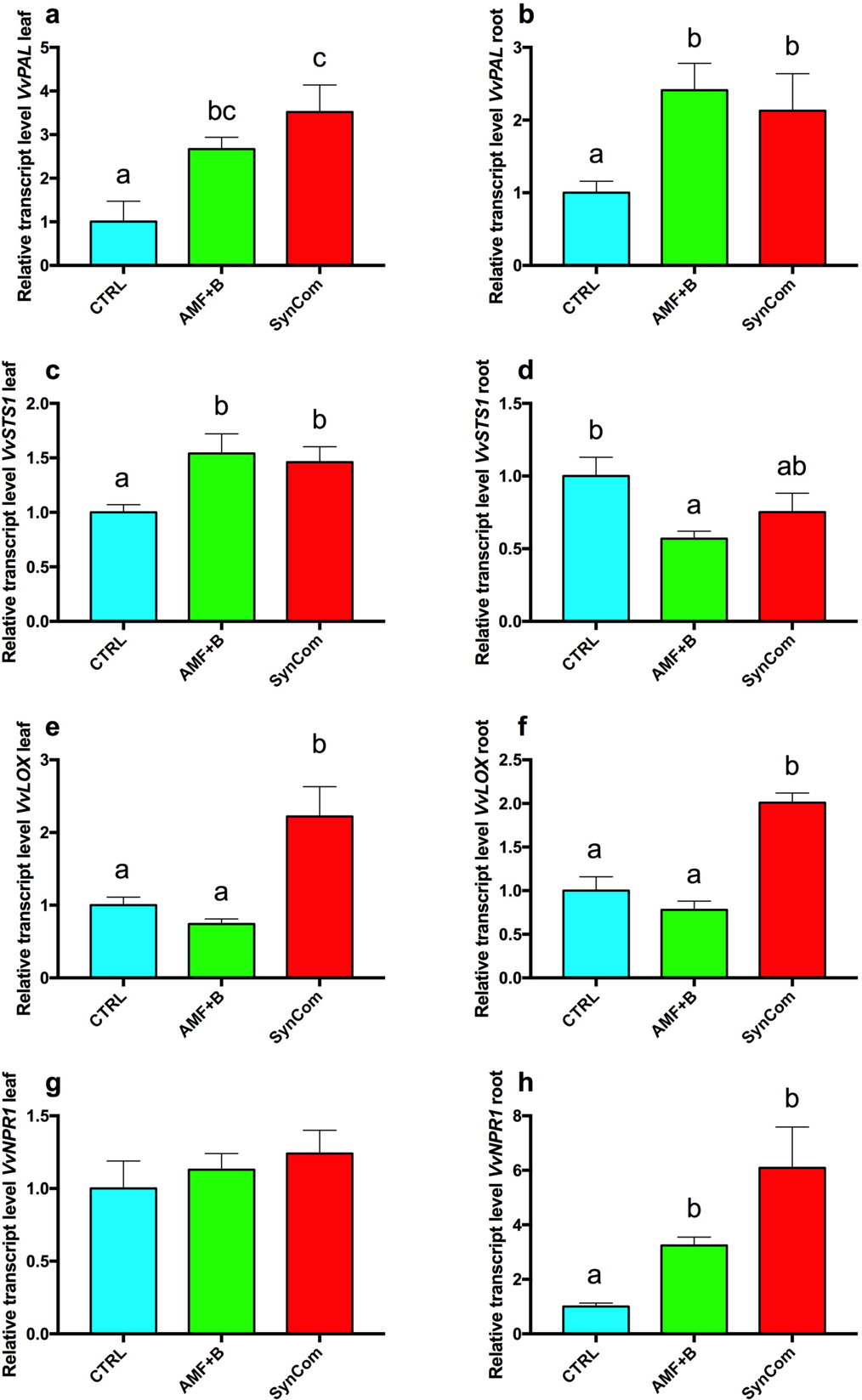
Expression changes of defence-related genes. **a,b)** Relative expression level of *VvPAL* in leaf and root tissues, respectively. **c,d)** Relative expression level of *VvSTS1* in leaf and root tissues, respectively. **e,f)** Relative expression level of *VvLOX* in leaf and root tissues, respectively. **g,h)** Relative expression level of *VvNPR1* in leaf and root tissues, respectively. Data are expressed as mean ± SD (n = 3). Different lowercase letters above bars indicate significant differences according to Tukey’s HSD test (*P* < *0.05*). CTRL, control plants; AMF+B, commercial AMF + Bacteria mixed inoculum-treated plants; SynCom, Synthetic Community-treated plants.

A second group of genes, associated with plant’s growth and physiological responses, were analysed to unravel further differences occurring among treatments. The expression of the ABA-biosynthetic gene *VvNCED3* (9-cis-epoxycarotenoid dioxygenase 3) was significantly down-regulated in the leaves of AMF+B- and SynCom-treated plants with respect to CTRL (Fig. S5a). For the same gene, a completely opposite transcriptional pattern was observed in the roots, with transcriptional levels that were almost double in all treated vines than those measured in the CTRL (Fig. S5b). *VvYUC3*, a key gene involved in the auxin synthesis pathway and encoding a flavin monooxygenase-like protein, also displayed different expression patterns based on treatment and tissue. Nevertheless, unlike what observed for *VvNCED3*, *VvYUC3* was exclusively overexpressed in the leaves of SynCom-treated vines (Fig. S5c), as in the roots it was significantly down-regulated in both treatments with respect to CTRL (Fig. S5d). Finally, *VvCHL*, a chlorophyllase-encoding gene, showed significantly higher transcriptional values in the leaves of SynCom-treated plants when compared with the analysis of samples collected from AMF+B and CTRL vines (Fig. S5e).

### Focus on SynCom-mediated whole transcriptome reprogramming in the leaf

In search of further experimental evidence strengthening the SynCom-mediated activation of systemic defence signalling routes, whole transcriptome changes were analysed by high-throughput sequencing (RNAseq) in the leaves collected from SynCom-treated plants. RNAseq data were first processed to identify those transcripts that were differentially expressed in the SynCom vs CTRL comparison, by applying a p-value adjusted with Benjamin-Hochberg ≤ 0.5% and setting a fold-change (log_2_ transformed FPKM values of the SynCom/CTRL ratio) threshold ≤ -1 or ≥ +1. Output of this analysis indicated that 147 out of the 29,970 annotated grapevine genes (V1 annotation of the PN40024 reference genome) were significantly differentially expressed (DEGs) in the SynCom vs CTRL comparison. Among the obtained DEGs, 142 were significantly upregulated in leaves of SynCom-treated plants (Table S8), whilst only 5 were down-regulated (Table S9). This finding suggested that, although the SynCom inoculation was successful in determining changes in the plant’s physiological performances (Fig. 5) and, at the biochemical level, in the accumulation of specific defensive metabolites (Fig. 6 and S4), the overall leaf transcriptomic alterations induced by this treatment were poor. Nevertheless, it deserves attention that almost all DEGs were overexpressed in SynCom leaves (Fig. 8a, Table S8). Moreover, a Gene Ontology enrichment analysis, performed on all DEGs, revealed that transport, response to endogenous stimulus, photosynthesis and energy metabolism were the overrepresented functional gene categories (Fig. 8b). Particularly, within the transport functional group, several transcripts encoding aquaporins and metal ion transporters were found upregulated. In the other enriched categories, such as response to stress and hormone metabolism and signalling, transcripts encoding lipoxygenases (*e.g.*, *VvLOX*), ethylene and auxin responsive factors, alpha expansins, monoxygenases (*e.g.*, *YUC3*), and stress related proteins (*e.g.*, *VvPAL*) were also overexpressed (Table S10). Cellular processes, signalling and homeostasis and energy metabolism also belonged to the upregulated functional categories, and specifically included genes encoding enzymes or proteins involved in cell wall metabolism (*e.g.*, cellulose synthase, arabinogalactan protein) and photosynthesis-related processes (*e.g.*, Ribulose 1,5-bisphosphate carboxylase, ferredoxin and photosystem I and II-related proteins). Conversely, three out of the five genes found significantly downregulated in the SynCom *vs* CTRL comparison were associated with hormone metabolism, and specifically with jasmonate-mediated signaling pathways (Table S10). Collectively, the analysis of RNAseq data thereby attested that, although the leaf transcriptome was poorly perturbed by SynCom inoculation, a basal activation of cell signalling and defence-related gene categories occurred, most likely supporting the systemic nature of specific defence responses in these plants.

**Figure 8.**
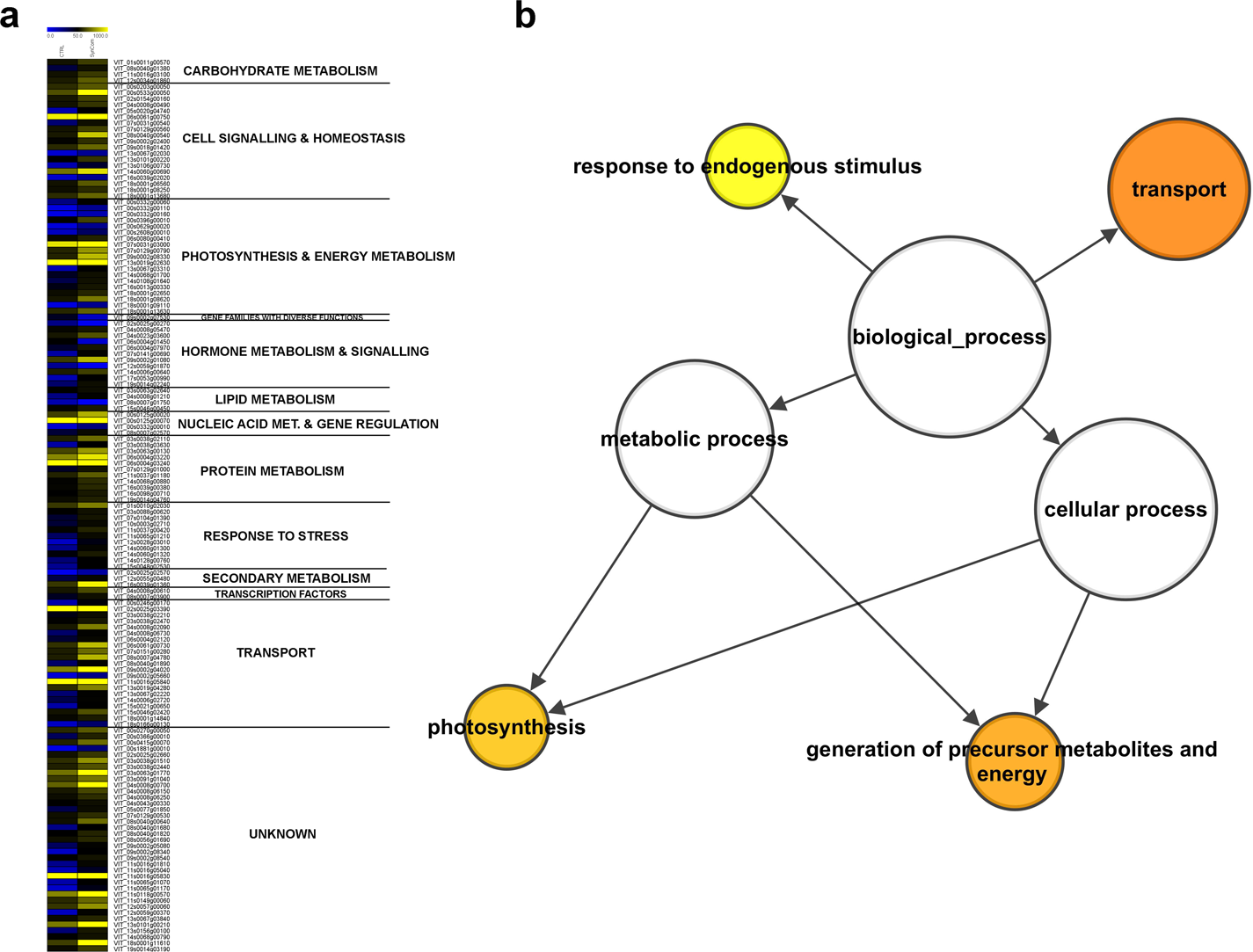
Analysis of whole leaf transcriptome. **a)** MeV heat map of differentially expressed transcripts. Colour scale of the heat map chart ranges from blue (low expression) to yellow (high expression, FPKM > 50); **b)** Significant enriched GO biological functional categories obtained by applying Cytoscape with the BINGO plug-in on the whole DEGs dataset and listed according to their enrichment p-value (*P < 0.05*).

## Discussion

To date, increasingly attention has been placed on the functions played by plant-associated microbes in response to environmental stresses (Marulanda et al., 2006; Sandrini et al., 2022). Since long-time scientists have reported beneficial effects conferred to plants by associated bacteria or fungi as a single strain or mixed to form consortia. The research focus was primarily posed on the use of soil microorganisms to boosting plant tolerance to drought conditions (Tufail et al., 2022). Nevertheless, the use of tailored SynComs to triggering plant immunity pathways against pathogens represents an approach still poorly explored (Li et al., 2021).

In search of experimental evidence supporting the application of SynComs to eliciting the plant’s defence system, a customized bacterial SynCom, formed by strains showing a marked *in vitro* biocontrol activity against *Vitis vinifera* fungal pathogens, was here formulated, considering isolates already identified as grapevine endophytes (Berendsen et al., 2018). The customized SynCom was inoculated in young grape cuttings grown in open field, and an integrated multidisciplinary approach was adopted to investigate its effect on plant physiological and endogenous defence responses, making comparison with the use of a commercial mixed inoculum and an uninoculated control.

### Biocontrol-based SynCom formulation

All bacterial strains were previously isolated from the inner woody tissue of the grape main trunk (*Vitis vinifera* cv Glera; Nerva et al. 2022a), ensuring in this way that the considered bacterial strains had the ability to live in strictly association with *Vitis vinifera* tissues. Outputs obtained from application of experimental SynComs in field conditions could be often contrasting (Ownley et al., 2003) and one of the reported main causes is the use of non-specific microbial cocktails exerting only mild effects on their plant hosts (Sandrini et al., 2022). The number of isolates forming SynComs greatly varied among research studies, from 3-4 to hundreds, as reviewed in (Sandrini et al., 2022). In the present survey, many isolates were proven to successfully limit the growth of different fungal pathogens, at least *in vitro*. Among those isolates, the seven most performing ones (two Actinobacteria and five Proteobacteria) were selected to build the grape customized SynCom. The reason for the use of such simplified SynCom relies on the fact that plants excrete in soil by roots a considerable amount of root exudates to attract microbes (Massalha et al., 2017) and the more isolates are applied, the greater is the competition among themselves (Li et al., 2021). To limit such effect, in our study a simplified SynCom was formulated using isolates that successfully confirmed their compatibility after a mutual exclusion test. Furthermore, the presence of Actinobacteria in the tailored SynCom is particularly relevant as their exploitation as sustainable agricultural tool has already been discussed (Viaene et al., 2016), although only a few species of this family were used as biocontrol agents (Bressan, 2003).

In the present study, the potential of bacterial endophytes as biocontrol agents in agriculture was mirrored by the fact that about 50% of the tested isolates exhibited a pathogen growth inhibition rate higher than 45% towards at least one of the four tested pathogens. Since no VOC activity was detected by means of septate-petri dish assay, such inhibition was probably due to diffusible compounds (Wan et al., 2008; Olivera et al., 2021). Although the main objective was the development of a customized SynCom formed by biocontrol agents able to trigger direct and indirect defence responses against grape pathogens, the seven isolated strains were also screened for some of the main PGP traits, such as nitrogen fixation, phosphate and starch solubilization or siderophore production, IAA production and ACC-deaminase activity. Additionally, the identified bacteria were also tested for tolerance to abiotic stress, *i.e.*, salt stress, which is becoming a frequent issue in viticulture due to both high fertilization rates and increasing incidence of drought events (Corwin, 2021). Among the seven PGP traits evaluated *in vitro*, each candidate isolate was found to be positive for at least two of the evaluated traits, highlighting its potential to improve plant growth and abiotic stress tolerance. However, such effects were not observed in field conditions, probably due to the occurrence of complex environmental interactions that could have masked them (Li et al., 2021).

### Plants inoculated with the customized SynCom showed growth-defence unbalanced responses in field with respect to AMF+B-treated vines

Although literature on the study and formulation of SynComs is available, only a few studies concern their application in glasshouse or experimental fields (Armanhi et al., 2021), and, at the best of our knowledge, no one was carried out in grapevine. As already stated, our survey was conceived to determine whether the application of synthetic communities (both customized or commercialized: SynCom or AMF+B, respectively) can represent a feasible approach in viticulture. In detail, the customized SynCom was inoculated on rooted cuttings before fielding them to foster the interaction establishment during the early developmental stages of rooted cuttings. Accordingly, Carlström et al. (2019) have recently demonstrated that community assembly is historically contingent and subjected to priority effects, so that the early timing of microbiome inoculation is essential to obtain a stable SynCom in planta. Additionally, adult plants are characterized by rich and complex microbiomes that remain largely unaffected by latecomers (Toju et al., 2018). It has been reported that individual strains of both Proteobacteria and Actinobacteria (both present in our SynCom) greatly affect the community structure as keystone species (Carlström et al., 2019). The metabarcoding data demonstrated that the species introduced with SynCom and AMF+B applications were still present in significantly high percentages several months after the inoculation (i.e., at the end of the season). The influence of plant-associated microorganisms on plant fitness has been already demonstrated (Yu et al., 2019). However, novel information on the effect of microbial inoculants on plant physiology is needed, and mainly when microorganisms are used in combination. To gain more insights into this subject, different physiological parameters were evaluated here. The main result was that, compared with CTRL, the two treatments (SynCom and AMF+B) differently modulated the vine’s growth-defence trade-off. Particularly, AMF+B-treated plants showed improved photosynthetic performances, suggesting that the synergic activity of AMF and rhizobacteria exerted a positive effect on plant fitness (Balestrini et al., 2020; Nerva et al., 2021a). Conversely, a negative impact on photosynthetic rates, ACE and iWUE was observed in SynCom-treated plants with respect to both AMF+B and CTRL ones. This finding was also consistent with changes in the expression profiles of *VvCHL*, a gene encoding a chlorophyllase (Chitarra et al., 2018), which was more transcribed in the leaves of SynCom-treated plants. Taken together, these data pointed to a growth imbalance phenomenon occurring in these plants, most likely caused by a SynCom-based establishment of specific defence responses.

To investigate this scenario more in depth, stress-related hormones were also analysed. Increased endogenous levels of ABA were found in roots of both SynCom- and AMF+B-inoculated plants. ABA is well known to play key roles in plants by improving stress tolerance and adaptation strategies to stressful factors (Egamberdieva et al., 2017). The observed patterns of ABA accumulation could suggest an impact on the plant’s ability to endure abiotic stress, making colonized plants more tolerant to climate alterations than CTRL vines (Sharp and LeNoble, 2002). Additionally, concentrations of IAA, a well-known growth marker (Teale et al., 2006), were not affected by treatments in leaves, while in roots they significantly increased but exclusively in the SynCom-inoculated plants. This finding could further imply that SynCom and AMF+B treatments differently shaped the plant growth and developmental pathways.

Bacterial communities were also reported to activate plant immunity mechanisms by priming the host to react against pathogen infections (Sandrini et al., 2022 and references therein). Stilbenes, such as resveratrol and viniferin, are the main defence-related metabolites synthesized in grapevine and displayed well documented antioxidant and antifungal properties, which are modulated by several factors, including the plant’s associated microbiota (Verhagen et al., 2010). Moreover, it is well documented that beneficial microbes can trigger diverse ISR-related pathways, most of which associated with the synthesis of stilbenes, including resveratrol and viniferin (Verhagen et al., 2010; Aziz et al., 2016; Nerva et al., 2021a). Our analysis showed that both AMF+B and SynCom treatments led to a sharp increase in *t*-resveratrol and viniferin amounts in roots and leaves, highlighting a microbe-dependent stimulation of defensive compound production (Li et al., 2021; Nerva et al., 2021a). These findings also strengthened the notion that in grapevine the primary mechanism for inducing systemic defence responses involves the production and accumulation of stilbenes. Besides stilbenoids, jasmonate is widely recognized as a key component of the ISR signalling in plants. An enhanced biosynthesis of MJ was exclusively noticed in the roots of plants treated with AMF+B and SynCom, apparently suggesting the establishment of localized rather than systemic defence responses in the treated plants.

To verify the impact of the two consortia application on the native microbial communities, a metabarcoding approach was adopted. This analysis allowed the characterisation of microbial populations associated with the roots three months after inoculation. AMF+B had a significant effect in reducing only the abundance of the *Neofusicoccum* and *Phaeoacremoniun* genera, whilst the impact of the SynCom treatment, besides stronger, was also evident on the *Phaeomoniella* genus. In this context, it should be considered that grape wood pathogens are often associated with severe symptoms only when their abundance reaches a certain threshold (Nerva et al., 2022a). Furthermore, to understand whether the observed effects were due to either direct antagonistic or indirect plant-mediated interactions (Guo et al., 2021), the network co-occurrence of the microbial dataset was analysed. It emerged that the main AMF+B effects were addressed to the stimulation of resveratrol accumulation (co-presence correlation between *Funelliformis* and *t*-resveratrol), which in turn was negatively correlated with the pathogen relative abundance. Interestingly, the bacterial members of the SynCom displayed both a direct and indirect antagonistic effect. It is worth noting that the introduced species belonged to genera that were negatively correlated with wood pathogens, but that had a positive correlation with *t*-resveratrol and, more importantly, with viniferin, one of the most important antifungal molecules in grapevine (Jeandet et al., 2002). Collectively, metabarcoding data indicated that the SynCom treatment was more efficient than AMF+B inoculation in reducing wood fungal pathogens, most likely due to a combination of direct antagonistic interactions (*e.g.,* pathogen abundance was negatively correlated to the abundance of introduced bacterial species) and indirect plant-mediated responses.

In summary, the integration of physiological, biochemical and metabarcoding analyses attested that SynCom-treated plants experienced improved defence responses accompanied by reduced photosynthetic performances. Such conditions suggested a shift of energy source allocation towards defence reactions. The AMF+B-inoculated vines mounted only a mild activation of endogenous defence signals and did not experience inhibition of photosynthetic rates, as reported previously (Augé et al., 2016; Chitarra et al., 2016; Goddard et al., 2021; Nerva et al., 2021a; Nerva et al., 2023).

### The analysis of stress-responsive and defence-related genes unveils molecular signals controlling target physiological and biochemical traits

Plants are known to finely tune their immune system during the interaction with beneficial microorganisms, regulating the expression of genes involved in different defence pathways (Alagna et al., 2020). Here, key defence-associated genes were analysed by means of qPCR. Among those, *VvSTS1* (a stilbene synthase gene) was overexpressed in leaves of both SynCom and AMF+B plants and downregulated in the roots of the same vines. Such opposite trend could likely rely on the high amount of both resveratrol and viniferin measured in roots, which could play a feedback negative regulation on the gene transcription. These data also confirmed a tissue-specific elicitation of plant’s immunity responses (Nerva et al., 2021a). Notably, a role for stilbenes in controlling accommodation of beneficial microorganisms in the roots and maintaining a homeostasis in the whole plant-associated microbial community has been reported as well (Liu et al., 2020).

At molecular level, the presence of treatment-dependent changes was also evaluated, looking at the expression of ISR- and SAR-related marker genes (*VvLOX* and *VvNPR1*, respectively), stress responsive and hormone-associated genes in root and leaf tissues. Although in AMF+B-treated vines ISR marker transcripts (*VvLOX*) were not induced, it is long-time known that AM plants can develop an enhanced defensive capacity against pathogens through the activation of the so-called ‘mycorrhiza-induced resistance’ (MIR) (Cameron et al., 2013), which in turn shares some features with the SAR and ISR paths (Bruisson et al., 2016; Goddard et al., 2021). Svenningsen et al. (2018) reported that AMF ecosystem services might be negatively affected by some rhizosphere bacterial groups, supporting the absence of *VvLOX* expression modulation in the AM-colonized plants.

Among defence-associated genes, the upregulation of *VvPAL* upon both treatments confirmed the elicitation of secondary metabolic pathways involved in preventing pathogen spread (Van Huylenbroeck et al., 1998; Oh et al., 2009; Giudice et al., 2022), consistently with the significant increase in the stilbenoid content in the same samples. Based on the notion that ABA is accumulated in response to different abiotic and biotic stresses (Ton et al., 2009; Egamberdieva et al., 2017), the expression of the ABA biosynthetic gene *VvNCED3* was also inspected. Similarly to ABA accumulation pattern, *VvNCED3* transcription was strongly activated in the roots of both AMF+B- and SynCom-treated plants, proving that ABA biosynthesis was enhanced upon the plant’s interaction with root-associated beneficial microbes (Chitarra et al., 2016). Martín-Rodríguez et al. (2016) reported that ABA metabolism is finely tuned in AM-colonized roots of tomato, but to our knowledge this is the first time that effects of SynCom application are positively correlated with increase in ABA concentration.

Furthermore, five out of the seven isolates forming the SynCom were IAA producers. Consequently, the plant auxin metabolic balance was also checked at the molecular level to verify the impact of the IAA released by the inoculated bacteria. Accordingly, a down-regulation of the auxin metabolic gene *VvYUC3* occurred in roots of both SynCom and AMF+B plants with respect to the CTRL. This data could likely suggest that the plants were using the IAA produced by the SynCom, thereby saving energy to be invested in other processes, such defensive ones. It was also reported that, by increasing the auxin pool available to the plants, bacterial IAA Producers (BIPs) positively affect the root system elongation and development, allowing water and nutrient uptake (Guerrieri et al., 2020). Additionally, a crosstalk between IAA production and induction of SA biosynthesis was noticed in IAA-primed wheat seeds (Iqbal and Ashraf, 2007). This could explain the high transcriptional rates of *VvNPR1*, a SAR marker gene, and mainly the high IAA amounts exclusive of the roots of SynCom-treated vines. Unlike bacteria, AMF have never reported as IAA producers, thereby supporting the reduced IAA content observed in AMF+B roots with respect to SynCom-inoculated samples.

### Leaf transcriptome profiling reveals the systemic nature of defence molecular signature in SynCom-treated plants

The findings discussed so far thus suggested that following SynCom application the endogenous plant’s defence machinery was promptly turned on. Such condition resulted in the activation of specific molecular (*e.g.*, exclusive *VvLOX* upregulation) and biochemical signals (*e.g.*, higher amounts of MJ and IAA), and also in peculiar physiological adjustments, as the reduction in assimilation rates. All those responses were either absent or moderately induced upon AMF+B treatment. These achievements also allowed us hypothesising that SynCom-promoted defence signalling routes operated systemically. To provide further evidence on this scenario, whole transcriptome reprogramming events were investigated in the leaf of SynCom-inoculated plants.

From a first analysis of sequencing data, it emerged that the leaf transcriptome was only poorly perturbed by the SynCom treatment. In fact, only 147 genes were found differentially expressed in the comparison with the transcriptome of CTRL leaves. Although we cannot exclude here a variability effect due to the intrinsic nature of samples taken from open field conditions, similar results were reported in studies conducted *in vitr*o or in controlled environment. Brotman et al., (2012) showed that *Arabidopsis thaliana* plants inoculated at the root level with a beneficial fungus had an improved resistance to the leaf pathogen *Pseudomonas syringae*. The authors proved that a systemic activation of the plant defence system already occurred in the inoculated plants before pathogen infection but without leading to a massive alteration of gene expression changes in the leaf, as only a few key defence transcripts were upregulated (Brotman et al., 2012). Consistently, Verhagen et al., (2004) had previously demonstrated that, following root colonization with the non-pathogenic bacteria *Pseudomonas fluorescens*, the onset of ISR signalling pathways observed in Arabidopsis leaves was not underlaid by an extensive transcriptome reprogramming.

Additionally in our study, grouping of differentially expressed transcripts into functional categories highlighted an enrichment in gene clusters specifically involved in defence metabolic pathways and regulation of stress response. This data, associated with the fact that almost all DEGs were overexpressed (being 142 out of 147 upregulated), further implied that the grapevine’s interaction with the customized SynCom channelled the plant metabolism towards the activation of defence mechanisms, hence shifting the growth-defence trade off. These results are in agreement with a recent study reporting a reduction in plant biomass and root growth and an overall low number of DEGs in the leaves of tomato plants inoculated with a microbial consortium (Prigigallo et al., 2023).

Together, the analysis of whole transcriptome changes occurring in the leaf of SynCom-inoculated plants confirmed that SynCom activity was able to orchestrate a specific reprogramming of genes involved in stress response, hormone metabolism and cell signalling. Additionally, these findings further proved that the SynCom-mediated trigger of defence metabolic pathways was not restricted to the root, and that a systemic activation of such responses was established resulting in a molecular signature at the leaf level.

#### Concluding remarks

In summary, since the new developed SynCom is formed by antagonistic bacteria with strong antibiosis activities against some fungal pathogens, they may, directly or indirectly, boost the plant immune system and the related defence responses. Such responses may hide their PGP effects observed *in vitro*, therefore shifting the growth-defence trade off towards defence responses.

Based on the above, the exploitation of synthetic communities selected for biocontrol activity could represent an interesting tool to manage the unbalanced growth-defence trade-off of modern genotypes (Nerva et al., 2022b). In this study, a simplified SynCom formed by seven grape bacterial isolates (firstly identified in a previous metatranscriptomic study by Nerva et al., 2022a) was formulated and inoculated in young cuttings. In addition to a non-inoculated control, the impact of the tailored SynCom on the physiological performances of field grown vines was compared with that of a commercial SynCom (formed by AMF and one rhizosphere bacterial species).

Results proved that the customized SynCom successfully boosted the vine’s defence machinery, while lowering photosynthetic performances, including carboxylation efficiency and water use efficiency due to an imbalance in the growth-defence trade off (Karasov et al., 2017). The adopted integration of physiological, biochemical and molecular data provides a clear picture of the responses occurring between the plant and its inhabiting microbiome, confirming that a holistic vision can shed light on the “dark side-effects” of SynCom application (Fig. 9).

**Figure 9.**
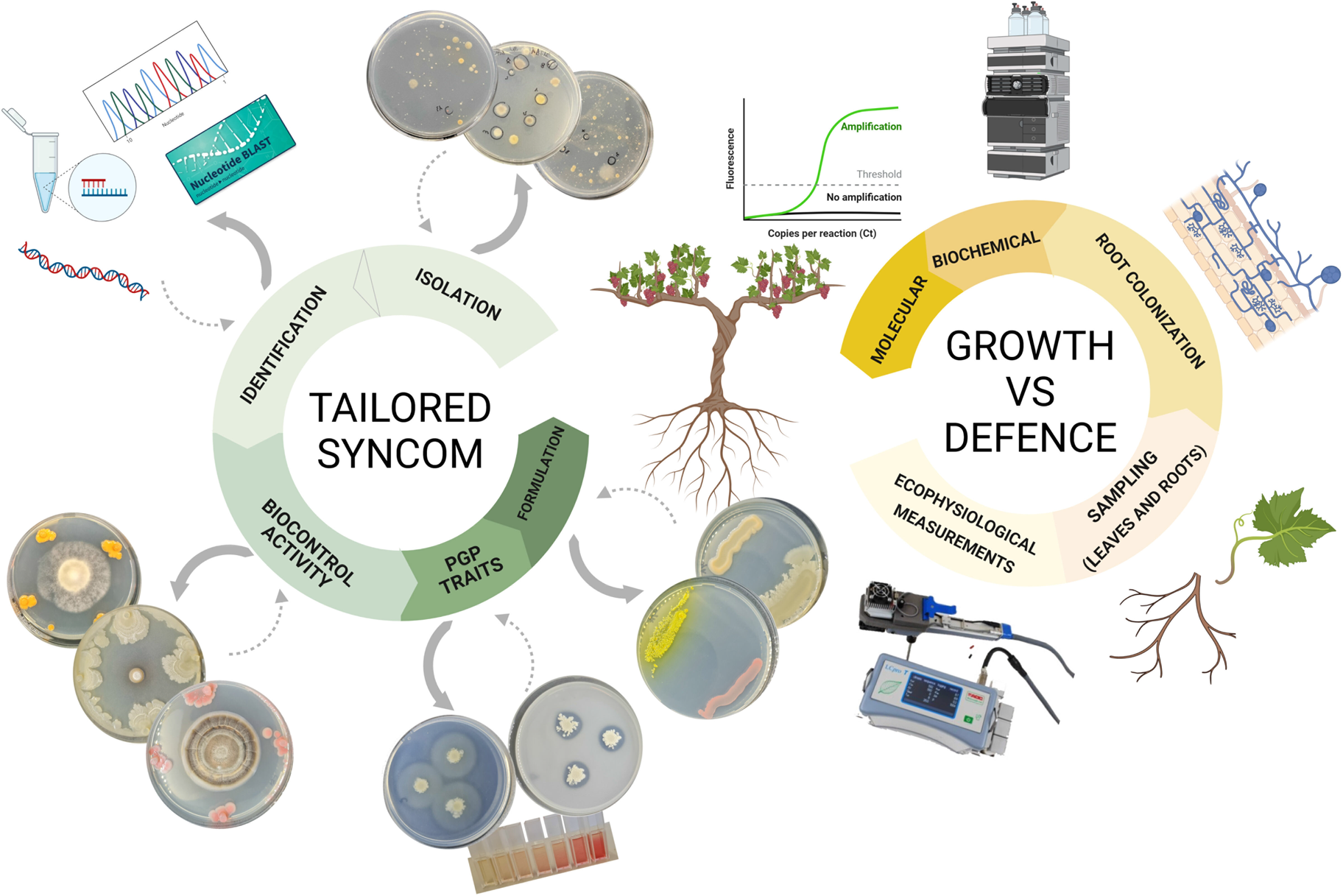
Overview of the adopted holistic approach. To test the SynCom developed in this work, several specific features of each isolate (*e.g.*, PGP traits, biocontrol activity, formulation characteristics, etc…) were considered and, once formulated, the holobiont responses were analyzed to decipher the impact on the plant trade-off features. The followed experimental strategy allowed observing physiological, biochemical and transcriptional rearrangements induced by the SynCom application.

Collectively, our achievements showed that the use of a similar *ad hoc* SynCom could be instrumental in those viticultural areas interested by high disease pressure, although negative effects on the plant photosynthetic performances potentially affecting grape production could occur. Nonetheless, the results obtained using the AMF+B inoculum suggest that the trigger of defence responses following the SynCom inoculation could be potentiated with the contemporaneous application of AMF isolates, leading to improved eco-physiological performances and defence reactions against pathogens in both aerial and underground parts of the plant.

Lastly, the development of tailored SynComs represents a very useful approach for research purposes aimed at dissecting the microbiome-host interaction mechanisms that are still poorly explored in field environment. Notably, building tailored SynComs (using both beneficial bacteria and fungi) for specific environmental settings represents a promising weapon to reducing the requirement of agrochemical and/or water inputs.

## Materials and methods

### Culture-dependent identification of bacteria isolates

To ensure the capability to stably colonize grape tissues, cultivable pure bacteria were previously isolated from the inner woody tissue of field-grown *Vitis vinifera* cv Glera plants as reported by (Nerva et al., 2022a) and stored in the CREA – Research Centre for Viticulture and Enology microbial bank (https://www.revine-prima2020.org/vimed) both as lyophilized and glycerol stocks (40% v/v) at -80°C. For Actinobacteria isolation a previously described method used: briefly, fresh wood tissue was grinded and resuspended in water amended with 0.1% SDS. The tissues with the water was vortexed for at least 45 minutes, diluted 1:5 with water and plated on sodium propionate medium (SPM) (Jiang et al., 2016). Fifteen days after plating, colonies were isolated using sterile needles and moving the bacterial colonies on CYA media. After total genomic DNA extractions, forty-four isolates were identified using the 16S sequence amplification (universal 27F 5’-AGAGTTTGATCCTGGCTCAG-3’ and 1392R 5’-GGTTACCTTGTTACGACTT-3’) and sequenced by Sanger method at BioFab Research srl (Italy). A search for similar sequences was conducted with the BLASTn tool (Basic Local Alignment Tool) on the GeneBank database, as reported in Table 1.

### *In vitro* antagonism screening against grape fungal pathogens and volatile effect

The ability of bacterial isolates to control *Botrytis cinerea* (bunch gray mold causal agent), *Phaeoacremonium minimun*, *Neofusicoccum parvum* (the latter two are key players in the esca syndrome) and *Guignardia bidwellii* (black rot causal agent) was evaluated *in vitro*. To perform biocontrol activity tests, each isolate was grown on solid media (CYA) for at least 5 days and then inoculated four times at the boundary of a 90mm Petri dish containing CYA media. The specific pathogen was inoculated 48 hours later as mycelial plug or as conidial suspension according to specific pathogen characteristics and then monitored for growth. Each combination of bacterial isolate and fungal pathogen was made in triplicate and the colony diameter measured twice for each biological replicate and each time point. For *B. cinerea* and *N. parvum* (both considered as fast-growing fungi) the biocontrol assay lasted 10 days at 28°C in dark conditions, for *P. minimum* and *G. bidwellii* (both considered slow-growing fungi) the assay lasted 20 days at 28°C in dark conditions. Inhibition percentages were calculated measuring the colony diameters on CYA of pathogens alone or in presence of bacterial isolates. The growth ratio was obtained by dividing the colony diameter on CYA with the bacteria over the colony diameter on CYA alone.

Additionally, when inhibition was observed, the potential antifungal activity of volatile organic compounds (VOCs) was evaluated according to (Oukala et al., 2021). In detail, the two room-plate method was used against the fungal pathogens reported above using at least three replicates for each bacterial isolate. After incubation at 28 °C, the percentage of mycelial growth inhibition was recorded and compared with their respective controls (Table 1).

The 7 best performing isolates were selected for the evaluation of PGP traits (see below) and for SynCom formulation.

### *In vitro* evaluation of plant growth promoting traits and compatibility test

*In vitro* compatibility test was performed in triplicate among the 7 selected isolates (Table 1, bold highlighted isolates) on CYA plates. Each bacterial strain, after overnight culture (500 µL, CYA media), was streaked onto solid media plates and co-cultured with each of the others. After ten days of incubation at 28°C, the bacterial growth was compared with a control plate where the isolate was cultured alone and, if present, inhibition effects were reported.

The selected isolates were also screened and evaluated for different PGP traits, such as production of indole acetic acid (IAA) (Guerrieri et al., 2020), ACC-deaminase activity (Li et al., 2011), siderophore production (Louden et al., 2011), N fixation (Geetha et al., 2014) using Jensen’s Nitrogen Free bacteria (JNFb) medium, phosphorus solubilization (Singh et al., 2020), starch hydrolyzation (Kokare et al., 2004) and salt stress resilience at diverse % of NaCl (0%, 1,5%, 3% w/v) (Gopalakrishnan et al., 2014) (Table 2). To evaluate IAA-production and ACC-deaminase activity, quantitative methods were adopted, whereas for siderophore production, phosphorus solubilization, starch solubilization and salinity resistance semi-quantitative methods were used. Finally, N fixation activity was assessed by qualitative method (presence or no presence of growth). In detail, regarding to phosphorus solubilization, bacterial ability was analysed measuring the halo zone diameter around the bacterial colonies: an isolate with a halo bigger than 0 mm up to 1 mm was considered as slight activity (+), a halo bigger than 1 mm up to 5 mm as medium activity (++) and a halo bigger than 5 mm as high activity (+++). Data for siderophore production was instead classified as follows: a zone of yellow halo appearance bigger than 0 mm up to 2 mm was evaluated as slight siderophore production, a zone of yellow halo appearance higher than 2 mm up to 5 mm as medium siderophore production and a zone of yellow bigger than 5 mm as good siderophore production. Moving to starch solubilization, data were classified as: a halo bigger than 0 cm up to 0.4 cm was considered as slight solubilization (+), a halo bigger than 0.4 cm up to 1 cm as medium solubilization (++) and a halo bigger than 1 cm as high solubilization (+++). Finally, for salinity resistance three different classes were set based on bacterial growth intensity: low growth (+), medium growth (++), good growth (+++).

### Plant inoculation and field experiments

Two hundred and thirty cutting vines of ‘Pinot gris’ cultivar grafted onto Kober 5BB and certified as ‘virus free’ were purchased from an Italian nursery (Vivai Cooperativi Rauscedo, Italy; http://www.vivairauscedo.com). Cuttings were treated as previously reported in (Nerva et al., 2021a) prior to plantation. Three treatments were compared in this study: i) non-treated control plants, CTRL; ii) SynCom-inoculated plants, SynCom; iii) SynCom-inoculated plants with a commercial consortium formed by different AMF species and a rhizosphere bacterial strain, AMF+B. The experiments were repeated twice, two independent rounds of cuttings inoculation and transplanting were performed for both SynComs (50+50 plants for each) while as CTRLs, 15 and 15 uninoculated plants were used for the first and second round of experiments.

As cited before, in this study we designed a 7-strain SynCom with isolates showing strong antagonistic activities previously isolated from grapevine woody tissues (see above) (Nerva et al., 2022a). The seven selected and compatible strains were grown in liquid CY media for 48 hours, then bacteria were collected using a centrifuge and resuspended in sterile water. Bacteria were counted and diluted to 10^9^ cells mL^-1^. The single bacterial suspensions were mixed to form the SynCom and inoculated in roots of one-year ‘Pinot gris’ cuttings using equal volume of each strain (∼ 10^8^ cells mL^-1^). In detail, prior to field planting, cuttings were maintained for 30 days in a plastic container filled with sterilized substrate (80% sand and 20% peat) supplemented with the formulated SynCom to a final concentration of 10^6^ cells mL^-1^ of substrate. For the AMF+B treatment, the grapevine cuttings were inoculated with a soluble powder-based commercial SynCom (MycoApply® DR formulation, Sumitomo Chemical Agro Europe SAS), formed by an AMF mixed inoculum (*Rhizophagus irregularis*, *Claroideoglomus luteum*, *Claroideoglomus etunicatum*, *Claroideoglomus claroideum* corresponding to 1% of the total inoculum as reported in the label) with and by a rhizospheric bacteria (*Bacillus coagulans*, 2.180.000 UFC g^-1^), following the manufacturer’s instruction (the inocula was resuspended in sterile water to use the same conditions of the formulated SynCom). As for the 7-strain SynCom, AMF+B-inoculated cuttings were maintained for 30 days in containers with steam sterilized substrate amended with the commercial inoculum. For CTRL plants, cuttings were prepared similarly and maintained on the same substrate for 30 days but without any microbiological inoculum.

Trials were carried out in a semi-controlled experimental field, specifically dedicated to this experiment, located at Cantine Rauscedo, Rauscedo, Italy (GPS coordinates: 46.054978N, 12.816345E). The about 3000 m^2^ of vineyard available for this study was composed of sandy loam soil (pH 7.3; available P 8.4 mg kg^-1^; organic matter 1.70%; cation exchange capacity 22.11 mew 100 g^-1^) that was not cultivated for three years prior to our experiments. After 60 days from field planting, leaf ecophysiological measurements and sampling of leaf and root tissues for molecular and biochemical analyses were performed. The collected samples were freeze-dried and stored at - 80°C until use. A part of the root apparatus was used to estimate the level of mycorrhiza colonization (*i.e.*, arbuscule abundance in the root system by morphological observation of thin roots fragments as previously described (Chitarra et al., 2016; Nerva et al., 2021a). Briefly, the root apparatus was stained with cotton blue and 1-cm long fragments were mounted on slides (generally 20 fragments for each slide) and the observations were performed on 3 or 5 slides (60 cm of the root system or 100 cm). Then, fungal colonization was quantified using the MYCOCALC software. The following parameters were considered: F% frequency of mycorrhiza in the root system, M% intensity of the mycorrhizal colonization in the root system, a% arbuscule abundance in mycorrhizal parts of root fragments, A% arbuscule abundance in the root system, v% vesicle abundance in mycorrhizal parts of root fragments.

Data obtained from the two independent experiments were collected and biological replicates were mediated and analyzed (see below).

### Leaf gas exchange measurements

Instantaneous measurements of net photosynthesis (Pn), stomatal conductance (g_s_), intercellular CO_2_ concentration (Ci), apparent carboxylation efficiency (ACE, calculated as the ratio between Pn and Ci) and intrinsic water use efficiency (iWUE, defined as the ratio between Pn and g_s_) were carried out on 6 randomly selected vines from each treatment for the two independent experiments. For each plant, three fully developed non-senescent leaves at the same physiological age (4^th^ to 5^th^ leaf from the shoot apex, n=36 per treatment) were measured using a portable infrared gas analyzer (ADC-LCi T system; Analytical Development Company, BioScientific Ltd., UK), as previously reported (Belfiore et al., 2021). The measurements were taken using ambient parameters as follows: light intensity ranged from 1.600 to 1.700 µmol photons m^-2^ s^-1^, ambient temperature ranged from 25 to 28°C, and CO_2_ concentration in the air ranged from 420 to 440 ppm.

### Targeted metabolite analysis

Leaf and root collected samples were used to determine abscisic acid (ABA), indole-acetic acid (IAA), *t*-resveratrol and viniferin concentrations using a high-performance liquid chromatographer (HPLC) (Nerva et al., 2021a; Nerva et al., 2021b). Briefly, since gas exchange data and trends were similar and comparable between the two independent experiments, for each treatment one biological replicate was formed by pooling at least four leaves collected in the median part of the canopy, taken from two randomly selected plants, one for each independent experiment. Thus, for each treatment and biological replicate, 100 mg of freeze-dried sample were aliquoted with 1 mL of extraction buffer (80% methanol-H_2_O, 8:2 v/v, with 0.1% v/v of acetic acid). The mixture was then sonicated in an ultrasonic bath for 1h at maximum intensity. After sonication, samples were centrifuged at maximum speed for 10 min at 4°C and filtrated using a 0.20 µm PTFE membrane filter (Chromafil^®^ Xtra PTFE-20/13, Macherey Nagel). The supernatant was analyzed by an HPLC apparatus, Agilent 1200 Infinity LC system model G4290B (Agilent, Waldbronn, Germany) equipped with gradient pump, auto-sampler and column oven set at 30°C. A C18 column (4.6 mm x 150 mm, 5 µm, XTerra^®^RP18) was used for the chromatographic separations. Original standards of ABA, IAA, *t*-resveratrol and *t*-viniferin (purity ≥ 98.5% and ≥99% for the latter two respectively, Merck KGaA, Darmstadt, Germany) were used to prepare the calibration curves and for the identification by comparing retention time and UV spectra. Analysis was run in reverse phase adopting an elution gradient method: eluent A was 0.1% formic acid in water and eluent B was acetonitrile; flow rate was fixed at 500 µL min^-1^. From 10% to 35% of B in 20 min, from 35% to 100% of B for 5 min, from 100% to 10% in 1 min and conditioning for 10 min. Twenty microliters were injected for each sample and at least three biological replicates (n=3) were run for each treatment.

To analyze changes in the accumulation of defense-related hormones (methyl salicylate, methyl jasmonate and jasmonic acid), the same lyophilized root and leaf samples were extracted as previously described (Huang et al., 2015). The extracts were then injected into an Agilent 6890N-5973i mass spectrometer. Chromatographic separation was performed using a Restek Rxi-5ms column (30m × 0.25mm × 0.25 µm) to ensure high-resolution and reliable quantification of the analytes (Restek Corporation, Bellefonte, PA, USA). The injection parameters were set at a temperature of 280°C and a volume of 2 µL, with a helium flow rate of 1.1 mL min^-1^. The Selected-Ion Monitoring (SIM) mode was utilized to enhance both the accuracy and sensitivity of the method. In contrast to Huang et al., specific ions were employed for the quantification of each analyte. For instance, salicylic acid was quantified at an m/z of 138 and identified through m/z values of 92 and 120. Methyl salicylate was quantified using m/z 152 and identified at m/z values of 120 and 92. Jasmonic acid was quantified at an m/z of 210 and identified at m/z values of 151 and 83. Lastly, methyl jasmonate was quantified using m/z 224 and identified through m/z values of 151 and 83. Electron ionization was conducted at 70 eV, with source and quadrupole temperatures optimized at 230°C and 150°C, respectively.

### RNA isolation, cDNA synthesis, qPCR analysis and RNA sequencing

The same root and leaf samples collected for metabolite analysis, were processed to isolate RNA starting from at least 50 mg of lyophilized tissue using the Spectrum™ Plant Total RNA Kit (Sigma-Aldrich) following manufacturer’s instructions (Nerva et al., 2021b). RNA concentrations were checked using a Nanodrop™ (Thermo Fisher Scientific) apparatus. Then, RNA samples were treated with DNase I (Thermo Fisher Scientific) following manufacturer’s instructions. The absence of DNA contamination was checked prior cDNA synthesis by quantitative real-time PCR (qPCR) using *VvCOX* grapevine specific primer (Table S2). After DNAse treatment, samples were subjected to cDNA synthesis using the High-Capacity cDNA Reverse Transcription Kit (Applied Biosystems, Thermo Fisher Scientific), starting from 500 µg of total RNA.

qPCR runs were performed in a final volume of 10 µL using SYBR^®^ green chemistry (Bio-Rad Laboratories Inc.) and 1:5 diluted cDNA as template. Reactions were performed in a Bio-Rad CFX96 instrument (Bio-Rad Laboratories Inc.) using the following conditions: denaturation phase at 95°C for 3 min, followed by 40 cycles at 95°C for 10 s and 60°C for 30 s. Each amplification was followed by a melting curve analysis (65-95°C) with a heating rate of 0.5°C every 5 s. All reactions were performed with at least two technical replicates. Transcript relative expression levels were calculated by the comparative cycle method using plant reference genes (cytochrome oxidase and ubiquitin, *i.e.*, *VvCOX* and *VvUBI*) for gene expression normalization. Oligonucleotide sequences are listed in Table S1. Gene expression data were calculated as expression ratio (relative quantity, RQ) to CTRL plants. For each analyzed gene, melting curve data were reported in Table S11.

RNA isolated from SynCom and CTRL leaf samples were submitted to library preparation and sequencing to gain information on the systemic molecular signature of SynCom inoculation. To proceed with RNAseq analysis, an average of 4 µg per sample were sent to Macrogen Inc. (South Korea), where cDNA libraries were built (TrueSeq total RNA sample kit, Illumina) and sequencing performed adopting the Illumina Novaseq technology with an average output of 40M paired-end reads (100 bp length) for each sample. Raw sequences were deposited in NCBI under the BioProject PRJNA1026525, BioSample SAMN37766153 and SAMN37766154 and SRA objects SRR26348056 and SRR26348055.

The Artificial Intelligence RNA-seq Software AIR (accessible at https://transcriptomics.cloud) was used to analyze RNA-seq data. AIR accepts as input file the raw next generation sequencing Illumina data (fastq format). RNA-seq data was uploaded to the cloud and validated to automatically pair forward and reverse files as well as to check its format and integrity. Quality analysis was assessed using FastQC. The forward analysis included quality trimming, Differential Gene Expression (DGE) followed by a Gene Ontology Enrichment Analysis (GOEA) using the V1 version of the grape PN40024 transcriptome (accessible at http://www.grapegenomics.com/) (Minio and Cantu, 2022). Once the analysis was launched, bad quality reads were removed using BBDuk by setting a minimum length of 35 bp and a minimum Phred-quality score of 25. Afterwards, high quality reads were mapped against the reference genome using STAR (Dobin et al., 2013) with the end-to-end alignment mode, and gene expression quantification was performed with featureCounts (Liao et al., 2014). Transcripts were annotated using the V1 version of the reference grapevine genome (Grimplet et al., 2012) and grouped into functional gene classes according to VitisNet GO. Heat maps of transcriptional profiles associated with specific functional categories were generated with TMeV 4.9 217 (http://www.tm4.org/mev.html), using as input the average expression value (FPKM) of the three 218 biological replicates. The BiNGO 3.0 plug-in tool in Cytoscape (v3.2, U.S. National Institute of General Medical Sciences (NIGMS), Bethesda, MD, USA) was used for running the GO enrichment analysis (Maere et al., 2021). Over-represented Plant GO slim categories were then identified using a hypergeometric test setting the significance threshold at 0.05. Differentially expressed genes (DEGs) were identified in a pairwise comparison (SynCom vs CTRL) using a p-value of 0.05% adjusted with the Benjamin-Hochberg method and setting the fold change (log_2_ transformed FPKM values of the SynCom/CTRL ratio) at ≥ +1 or ≤ -1.

### Root DNA isolation and metabarcoding analysis

Starting from the root powder previously obtained, about 50 mg were used to extract DNA following the manufacturer’s instruction of the plant/fungi DNA isolation kit (Norgen Biotech Corp., Thorold, ON, Canada), as previously reported (Nerva et al., 2021c). For each treatment, four biological replicates were extracted by randomly selecting plants from the two experimental replications. Total DNA was quantified using a NanoDrop One spectrophotometer (Thermo Fisher Scientific, Waltham, MA, USA), and DNA integrity was inspected running the extracted samples on a 1% agarose electrophoretic gel. While preparing the samples for sequencing, a further quantification was performed using a Qubit 4 Fluorometer (Thermo Fisher Scientific, Waltham, MA, USA) to compare and integrate the previous quantifications.

To analyze the root-associated bacterial community the V3-V4 hypervariable region of the 16S rRNA gene was targeted by the universal primers 319F (5′-CCTACGGGNGGCWGCAG-3′) and 806R (5′-GACTACHVGGGTATCTAATCC-3′). For the inspection of the root-associated fungal community the ITS2 region of the rRNA gene was analyzed using the primers ITS3 (5’-GCATCGATGAAGAACGCAGC-3’) and ITS4 (5’-TCCTCCGCTTATTGATATGC-3’). The ad-hoc designed peptide nucleotide acid (PNA) blocker oligos (Kaneka Eurogentec S.A., Belgium) for *V. vinifera* were used to inhibit plant material amplification at plant mitochondrial and chloroplast 16S rRNA genes (mitochondrial and plastidial) and at plant 5.8S nuclear rRNA (Lundberg et al., 2013; Cregger et al., 2018; Nerva et al., 2021a). The obtained metabarcoding sequences were deposited in NCBI database under the BioProject PRJNA1026525, BioSample SAMN37748419-SAMN37748424 and SRA objects SRR26363391-SRR26363395. A first strict quality control on raw data was performed with FLASh (Magoč and Salzberg, 2011) and then processed with two independent Qiime2 (Bolyen et al., 2019) pipelines. To analyze the bacterial community, the 16S sequences were subjected to quality filtering with DADA2 (Callahan et al., 2016) to perform chimera removal, error-correction and sequence variant calling with reads truncated at 260 bp and displaying a quality score above 20. Feature sequences were summarized and annotated using the RDP classifier (Cole et al., 2014) trained to the full length 16S database retrieved from the curated SILVA database (v138) (Quast et al., 2012). For graphic representation, only genera with an average relative abundance higher than the settled threshold (2%) were retained with the exception for the genera recalling the inoculated isolates.

To analyze the fungal community through ITS2 sequences, reads were first filtered to identify the start and stop sites for the ITS region using the hidden Markov models (HMM) (Rivers et al., 2018). Briefly, the software allows to distinguish true sequences from sequencing errors, filtering out reads with errors or reads without ITS sequences. To distinguish true sequences from those containing errors, sequences have been sorted by abundance and then clustered in a greedy fashion at a threshold percentage of identity (97%). Trimmed sequences were analyzed with DADA2 and sequence variants were taxonomically classified through the UNITE database (Abarenkov et al., 2010). For graphic representation, only genera with an average relative abundance higher than the settled threshold (1%) were retained with the exception for genera of inoculated mycorrhizal fungi. Co-occurrence Network interference (CoNet v1.1.1.beta) (Faust and Raes, 2012; Faust and Raes, 2016) was employed to identify significant co-occurrence patterns among the bacterial communities. To such aim, the ASV table data was imported into Cytoscape (v3.9.1) (Shannon et al., 2003) through the CoNet app. The top 100 edges with the highest positive and negative values were selected and combined using the mean value through the union approach. Multi-edge scores were then shuffled row-wise at 100 permutations (for the randomization). The brown method (Wiese et al., 2004) was utilized to merge node pairs, which were assigned via the p-values of the multi-edges. Unstable edges were removed, and a significance threshold of *P* < 0.05 was applied to determine the q-value (the corrected significance value). The edges were coloured via their positive (co-presence; green) and negative (co-exclusion; red) association.

### Statistics

Data were analyzed by analysis of variance (ANOVA). When ANOVA was significant, mean separation was performed according to Tukey’s HSD test at a probability level of *P* ≤ 0.05. Standard deviation (SD) or error (SE) of all means was calculated. The SPSS statistical software package (version 22) was used to run statistical analyses.

Microbial data were analyzed using the qiime2-implemented functions (Kruskal–Wallis test, Permanova). Ecological indices (i.e., NMDS and PERMANOVA) were calculated in Past4.04. RNAseq data were analyzed using dedicated pipelines in Artificial Intelligence RNA-seq Software AIR (accessible at https://transcriptomics.cloud).

## Supplemental data

The following materials are available in the online version of this article.

**Figure S1.** Co-colture assay. To test the ability of the selected isolates to share the same environmental niche, we grew them on the same culture media (CYA) for at least one week in order to observe eventual presence of inhibition effects.

**Figure S2.** Non-metric multidimensional scaling (NMDS) and PERMANOVA analyses on bacterial and fungal communities. NMDS and PERMANOVA were calculated for both bacterial a) and fungal b) communities considering CTRL, AMF+B and SynCom treatments.

**Figure S3.** Network co-occurrence analysis for each treatment. Network analysis was performed by using as input the microbial abundance data (bacterial and fungi combined) and the biochemical measurements of CTRL (a), AMF+B (b) and SynCom (c). Green lines highlight copresence interactions, while red lines indicate mutual exclusion. Each node represents a taxonomic group.

**Figure S4.** Concentration of methyl-salicylate (MS), methyl-jasmonate (MJ) and jasmonic acid (JA) in leaf (a) and root (b) tissues.

**Figure S5.** Expression changes of photosynthesis and hormones-related genes. **a)** Relative expression level of *VvChl* in leaf. **b,c)** Relative expression level of *VvNCED3* in root and leaf tissues, respectively. **d,e)** Relative expression level of *VvYUC3* in leaf and root tissues, respectively. Data are expressed as mean ± SD (n = 3). Different lowercase letters above bars indicate significant differences according to Tukey’s HSD test (*P* < 0.05). CTRL, control plants; AMF+B, commercial AMF + Bacteria mixed inoculum-treated plants; SynCom, Synthetic Community-treated plants.

**Supplementary Table S1.** Evaluation of plant growth promoting (PGP) traits of selected bacteria constituting the SynCom. Symbol – indicates negative result and symbol + positive result. Positive results have been ranked with an intensity scale: +, slight activity; ++ medium activity; +++ good activity. Data for ACC-deaminase activity are expressed as ACC concentration remaining in the DF-ACC medium containing 3 mM ACC after incubation of each ACC-utilizing bacterial isolate for 24h. Data for IAA-production are expressed as IAA concentrations (µg/mL-1) after incubation of each isolate for 48h in Luria Bertani broth. Data for siderophore production: measurement (mm) of the yellow halo zone around the bacterial colonies on CAS agar plate. Data of N fixation: +, growth capability on Nfb agar medium. Data for P solubilization and starch hydrolysis: measurement of halo diameter (zone of clearance) around the bacterial colonies. Data related to salinity tolerance: intensity of bacterial growth on agar plate containing different NaCl concentrations. Data are the mean values of three replicates ± SD for each isolate.

**Supplementary Table S2.** List of the oligonucleotides used in this study for qPCR analysis of candidate gene expression. For each primer pair, the gene annotation, gene ID (GGDB 12X V1), amplicon length (bp), annealing temperature (Ta, °C).

**Supplementary Table S3**. Number of retained sequences after chimera removal and taxonomical assignment.

**Supplementary Table S4**. Shannon diversity index calculated for replicate in each group. Average, standard deviation and significance of pairwise comparison are reported.

**Supplementary Table S5**. Summary of the 16S data for each treatment. Average, standard deviation and p value of pairwise comparisons are reported.

**Supplementary Table S6**. Summary of the ITS data for each treatment. Average, standard deviation and p value of pairwise comparisons are reported.

**Supplementary Table S7**. Network analysis considering both the 16S and ITS interactions together with biochemical compounds. a) CTRL plants, b) AMF+B plants, c) SynCom plants, d) analysis on the whole dataset (CTRL, AMF+B and SynCom).

**Supplementary Table S8**. List of significantly upregulated genes in the comparison between SynCom-treated (SynCom) and reference (CTRL) plants.

**Supplementary Table S9**. List of significantly downregulated genes in the comparison between SynCom-treated (SynCom) and reference (CTRL) plants.

**Supplementary Table S10**. Distribution of functional gene categories. Functional categories of genes differentially expressed in the pairwise comparison between treated (SynCom) and control untreated (CTRL) plants.

**Supplementary Table S11.** Summary data on melting curves.

## Acknowledgments

The authors are grateful to Dr. Giancarlo Babbo formerly employed in Sumitomo Chem Italia for kindly providing MycoApply product used in this study.

## References

1. Abarenkov K, Henrik Nilsson R, Larsson K, Alexander IJ, Eberhardt U, Erland S, Høiland K, Kjøller R, Larsson E, Pennanen T (2010) The UNITE database for molecular identification of fungi–recent updates and future perspectives. New Phytol 186: 281–285

2. Alagna F, Balestrini R, Chitarra W, Marsico A, Nerva L (2020) Getting ready with the priming: Innovative weapons against biotic and abiotic crop enemies in a global changing scenario. Priming-Mediat. Stress Cross-Stress Toler. Crop Plants. Elsevier, pp 35–56

3. Armanhi JSL, de Souza RSC, Biazotti BB, Yassitepe JE de CT, Arruda P (2021) Modulating drought stress response of maize by a synthetic bacterial community. Front Microbiol 12: 747541

4. Armijo G, Schlechter R, Agurto M, Muñoz D, Nuñez C, Arce-Johnson P (2016) Grapevine pathogenic microorganisms: understanding infection strategies and host response scenarios. Front Plant Sci 7: 382

5. Augé RM, Toler HD, Saxton AM (2016) Mycorrhizal stimulation of leaf gas exchange in relation to root colonization, shoot size, leaf phosphorus and nitrogen: a quantitative analysis of the literature using meta-regression. Front Plant Sci 7: 1084

6. Aziz A, Verhagen B, Magnin-Robert M, Couderchet M, Clément C, Jeandet P, Trotel-Aziz P (2016) Effectiveness of beneficial bacteria to promote systemic resistance of grapevine to gray mold as related to phytoalexin production in vineyards. Plant Soil 405: 141–153

7. Balestrini R, Brunetti C, Chitarra W, Nerva L (2020) Photosynthetic Traits and Nitrogen Uptake in Crops: Which Is the Role of Arbuscular Mycorrhizal Fungi? Plants 9: 1105

8. Balestrini R, Salvioli A, Dal Molin A, Novero M, Gabelli G, Paparelli E, Marroni F, Bonfante P (2017) Impact of an arbuscular mycorrhizal fungus versus a mixed microbial inoculum on the transcriptome reprogramming of grapevine roots. Mycorrhiza 27: 417–430

9. Belfiore N, Nerva L, Fasolini R, Gaiotti F, Lovat L, Chitarra W (2021) Leaf gas exchange and abscisic acid in leaves of Glera grape variety during drought and recovery. Theor Exp Plant Physiol

10. Berendsen RL, Vismans G, Yu K, Song Y, de Jonge R, Burgman WP, Burmølle M, Herschend J, Bakker PA, Pieterse CM (2018) Disease-induced assemblage of a plant-beneficial bacterial consortium. ISME J 12: 1496–1507

11. Bolyen E, Rideout JR, Dillon MR, Bokulich NA, Abnet CC, Al-Ghalith GA, Alexander H, Alm EJ, Arumugam M, Asnicar F (2019) Reproducible, interactive, scalable and extensible microbiome data science using QIIME 2. Nat Biotechnol 37: 852–857

12. Bressan W (2003) Biological control of maize seed pathogenic fungi by use of actinomycetes. BioControl 48: 233–240

13. Brotman Y, Lisec J, Meret M, Chet I, Willmitzer L, Viterbo A (2012) Transcript and metabolite analysis of the Trichoderma-induced systemic resistance response to *Pseudomonas syringae* in *Arabidopsis thaliana*. Microbiology 158: 139–146

14. Bruisson S, Maillot P, Schellenbaum P, Walter B, Gindro K, Deglène-Benbrahim L (2016) Arbuscular mycorrhizal symbiosis stimulates key genes of the phenylpropanoid biosynthesis and stilbenoid production in grapevine leaves in response to downy mildew and grey mould infection. Phytochemistry 131: 92–99

15. Burketova L, Trda L, Ott PG, Valentova O (2015) Bio-based resistance inducers for sustainable plant protection against pathogens. Biotechnol Adv 33: 994–1004

16. Callahan BJ, McMurdie PJ, Rosen MJ, Han AW, Johnson AJA, Holmes SP (2016) DADA2: high-resolution sample inference from Illumina amplicon data. Nat Methods 13: 581–583

17. Cameron DD, Neal AL, van Wees SC, Ton J (2013) Mycorrhiza-induced resistance: more than the sum of its parts? Trends Plant Sci 18: 539–545

18. Carlström CI, Field CM, Bortfeld-Miller M, Müller B, Sunagawa S, Vorholt JA (2019) Synthetic microbiota reveal priority effects and keystone strains in the *Arabidopsis* phyllosphere. Nat Ecol Evol 3: 1445–1454

19. Carrión VJ, Perez-Jaramillo J, Cordovez V, Tracanna V, De Hollander M, Ruiz-Buck D, Mendes LW, van Ijcken WF, Gomez-Exposito R, Elsayed SS (2019) Pathogen-induced activation of disease-suppressive functions in the endophytic root microbiome. Science 366: 606–612

20. Chitarra W, Cuozzo D, Ferrandino A, Secchi F, Palmano S, Perrone I, Boccacci P, Pagliarani C, Gribaudo I, Mannini F (2018) Dissecting interplays between *Vitis vinifera* L. and *grapevine virus B* (GVB) under field conditions. Mol Plant Pathol 19: 2651–2666

21. Chitarra W, Pagliarani C, Maserti B, Lumini E, Siciliano I, Cascone P, Schubert A, Gambino G, Balestrini R, Guerrieri E (2016) Insights on the impact of arbuscular mycorrhizal symbiosis on tomato tolerance to water stress. Plant Physiol 171: 1009–1023

22. Chitarra W, Siciliano I, Ferrocino I, Gullino ML, Garibaldi A (2015) Effect of elevated atmospheric CO2 and temperature on the disease severity of rocket plants caused by Fusarium wilt under phytotron conditions. PloS One 10: e0140769

23. Cole JR, Wang Q, Fish JA, Chai B, McGarrell DM, Sun Y, Brown CT, Porras-Alfaro A, Kuske CR, Tiedje JM (2014) Ribosomal Database Project: data and tools for high throughput rRNA analysis. Nucleic Acids Res 42: D633–D642

24. Corwin DL (2021) Climate change impacts on soil salinity in agricultural areas. Eur J Soil Sci 72: 842–862

25. Cregger M, Veach A, Yang Z, Crouch M, Vilgalys R, Tuskan G, Schadt C (2018) The *Populus* holobiont: dissecting the effects of plant niches and genotype on the microbiome. Microbiome 6: 1–14

26. Dastogeer KMG, Chakraborty A, Sarker MSA, Akter MA (2020) Roles of fungal endophytes and viruses in mediating drought stress tolerance in plants. Int J Agric Biol 24: 1497–1512

27. Dobin A, Davis CA, Schlesinger F, Drenkow J, Zaleski C, Jha S, Batut P, Chaisson M, Gingeras TR (2013) STAR: ultrafast universal RNA-seq aligner. Bioinformatics 29: 15–21

28. Egamberdieva D, Wirth SJ, Alqarawi AA, Abd_Allah EF, Hashem A (2017) Phytohormones and beneficial microbes: essential components for plants to balance stress and fitness. Front Microbiol 8: 2104

29. Faust K, Raes J (2012) Microbial interactions: from networks to models. Nat Rev Microbiol 10: 538–550

30. Faust K, Raes J (2016) CoNet app: inference of biological association networks using Cytoscape. F1000Research 5

31. Geetha K, Venkatesham E, Hindumathi A, Bhadraiah B (2014) Isolation, screening and characterization of plant growth promoting bacteria and their effect on *Vigna Radita* (L.) R. Wilczek. Int J Curr Microbiol Appl Sci 3: 799–899

32. Giudice G, Moffa L, Niero M, Duso C, Sandrini M, Vazzoler LF, Luison M, Pasini E, Chitarra W, Nerva L (2022) Novel sustainable strategies to control *Plasmopara viticola* in grapevine unveil new insights on priming responses and arthropods ecology. Pest Manag Sci 78: 2342–2356

33. Giudice G, Moffa L, Varotto S, Cardone MF, Bergamini C, De Lorenzis G, Velasco R, Nerva L, Chitarra W (2021) Novel and emerging biotechnological crop protection approaches. Plant Biotechnol J 19: 1495–1510

34. Goddard M-L, Belval L, Martin IR, Roth L, Laloue H, Deglène-Benbrahim L, Valat L, Bertsch C, Chong J (2021) Arbuscular mycorrhizal symbiosis triggers major changes in primary metabolism together with modification of defense responses and signaling in both roots and leaves of *Vitis vinifera*. Front Plant Sci 12: 721614

35. Gopalakrishnan S, Vadlamudi S, Bandikinda P, Sathya A, Vijayabharathi R, Rupela O, Kudapa H, Katta K, Varshney RK (2014) Evaluation of *Streptomyces* strains isolated from herbal vermicompost for their plant growth-promotion traits in rice. Microbiol Res 169: 40–48

36. Grimplet J, Van Hemert J, Carbonell-Bejerano P, Díaz-Riquelme J, Dickerson J, Fennell A, Pezzotti M, Martínez-Zapater JM (2012) Comparative analysis of grapevine whole-genome gene predictions, functional annotation, categorization and integration of the predicted gene sequences. BMC Res Notes 5: 213

37. Guerrieri MC, Fanfoni E, Fiorini A, Trevisan M, Puglisi E (2020) Isolation and screening of extracellular PGPR from the rhizosphere of tomato plants after long-term reduced tillage and cover crops. Plants 9: 668

38. Guo Y, Luo H, Wang L, Xu M, Wan Y, Chou M, Shi P, Wei G (2021) Multifunctionality and microbial communities in agricultural soils regulate the dynamics of a soil-borne pathogen. Plant Soil 461: 309–322

39. Huang Z, Wang Z, Shi B, Wei D, Chen J, Wang S, Gao B (2015) Simultaneous determination of salicylic acid, jasmonic acid, methyl salicylate, and methyl jasmonate from *Ulmus pumila* leaves by GC-MS. Int J Anal Chem 2015

40. Iqbal M, Ashraf M (2007) Seed treatment with auxins modulates growth and ion partitioning in salt-stressed wheat plants. J Integr Plant Biol 49: 1003–1015

41. Jeandet P, Douillet-Breuil A-C, Bessis R, Debord S, Sbaghi M, Adrian M (2002) Phytoalexins from the Vitaceae: biosynthesis, phytoalexin gene expression in transgenic plants, antifungal activity, and metabolism. J Agric Food Chem 50: 2731–2741

42. Jiang Y, Li Q, Chen X, Jiang C (2016) Isolation and cultivation methods of Actinobacteria. Actinobacteria–Basics Biotechnol Appl 39–57

43. Karasov TL, Chae E, Herman JJ, Bergelson J (2017) Mechanisms to mitigate the trade-off between growth and defense. Plant Cell 29: 666–680

44. Kokare C, Mahadik K, Kadam S, Chopade B (2004) Isolation, characterization and antimicrobial activity of marine halophilic *Actinopolyspora* species AH1 from the west coast of India. Curr Sci 593–597

45. Lebeis SL (2015) Greater than the sum of their parts: characterizing plant microbiomes at the community-level. Curr Opin Plant Biol 24: 82–86

46. Li Z, Bai X, Jiao S, Li Y, Li P, Yang Y, Zhang H, Wei G (2021) A simplified synthetic community rescues *Astragalus mongholicus* from root rot disease by activating plant-induced systemic resistance. Microbiome 9: 1–20

47. Li Z, Chang S, Lin L, Li Y, An Q (2011) A colorimetric assay of 1-aminocyclopropane-1-carboxylate (ACC) based on ninhydrin reaction for rapid screening of bacteria containing ACC deaminase. Lett Appl Microbiol 53: 178–185

48. Liao Y, Smyth GK, Shi W (2014) featureCounts: an efficient general purpose program for assigning sequence reads to genomic features. Bioinformatics 30: 923–930

49. Liu H, Brettell LE, Qiu Z, Singh BK (2020) Microbiome-mediated stress resistance in plants. Trends Plant Sci 25: 733–743

50. Liu Y-X, Qin Y, Bai Y (2019) Reductionist synthetic community approaches in root microbiome research. Curr Opin Microbiol 49: 97–102

51. Louden BC, Haarmann D, Lynne AM (2011) Use of blue agar CAS assay for siderophore detection. J Microbiol Biol Educ 12: 51–53

52. Lundberg DS, Yourstone S, Mieczkowski P, Jones CD, Dangl JL (2013) Practical innovations for high-throughput amplicon sequencing. Nat Methods 10: 999–1002

53. Maere S, Heymans K, Kuiper M (21) June 2005, posting date. BiNGO Cytoscape Plugin Assess Overrepresentation Gene Ontol. Categ. Biol. Netw. Bioinforma. 21: 3448–3449

54. Magoč T, Salzberg SL (2011) FLASH: fast length adjustment of short reads to improve genome assemblies. Bioinformatics 27: 2957–2963

55. Sandrini M, Moffa L, Velasco R, Balestrini R, Chitarra W, Nerva L (2022) Microbe-assisted crop improvement: a sustainable weapon to restore holobiont functionality and resilience. Hortic. Res. 9: uhac160

56. Martín-Rodríguez JA, Huertas R, Ho-Plágaro T, Ocampo JA, Turečková V, Tarkowská D, Ludwig-Müller J, García-Garrido JM (2016) Gibberellin–abscisic acid balances during arbuscular mycorrhiza formation in tomato. Front Plant Sci 7: 1273

57. Marulanda A, Barea J, Azcón R (2006) An indigenous drought-tolerant strain of *Glomus intraradices* associated with a native bacterium improves water transport and root development in *Retama sphaerocarpa*. Microb Ecol 52: 670–678

58. Massalha H, Korenblum E, Tholl D, Aharoni A (2017) Small molecules below-ground: the role of specialized metabolites in the rhizosphere. Plant J 90: 788–807

59. Minio A, Cantu D (2022) Grapegenomics. com: a web portal with genomic data and analysis tools for wild and cultivated grapevines.

60. Nerva L, Balestrini R, Chitarra W (2023) From Plant Nursery to Field: Persistence of Mycorrhizal Symbiosis Balancing Effects on Growth-Defence Tradeoffs Mediated by Rootstock. Agronomy 13: 229

61. Nerva L, Garcia J, Favaretto F, Giudice G, Moffa L, Sandrini M, Cantu D, Zanzotto A, Gardiman M, Velasco R (2022a) The hidden world within plants: metatranscriptomics unveils the complexity of wood microbiomes. J. Exp. Bot. 73: 2682–2697

62. Nerva L, Giudice G, Quiroga G, Belfiore N, Lovat L, Perria R, Volpe MG, Moffa L, Sandrini M, Gaiotti F (2021a) Mycorrhizal symbiosis balances rootstock-mediated growth-defence tradeoffs. Biol Fertil Soils 58: 17–34

63. Nerva L, Guaschino M, Pagliarani C, De Rosso M, Lovisolo C, Chitarra W (2021b) Spray induced gene silencing targeting a *glutathione S-transferase* gene improves resilience to drought in grapevine. Plant Cell Environ. 45: 347–361

64. Nerva L, Moffa L, Giudice G, Giorgianni A, Tomasi D, Chitarra W (2021c) Microscale analysis of soil characteristics and microbiomes reveals potential impacts on plants and fruit: vineyard as a model case study. Plant Soil 462: 525–541

65. Nerva L, Pagliarani C, Pugliese M, Monchiero M, Gonthier S, Gullino ML, Gambino G, Chitarra W (2019) Grapevine Phyllosphere Community Analysis in Response to Elicitor Application against Powdery Mildew. Microorganisms 7: 662

66. Nerva L, Sandrini M, Moffa L, Velasco R, Balestrini R, Chitarra W (2022b) Breeding toward improved ecological plant–microbiome interactions. Trends Plant Sci.

67. Oh M-M, Trick HN, Rajashekar C (2009) Secondary metabolism and antioxidants are involved in environmental adaptation and stress tolerance in lettuce. J Plant Physiol 166: 180–191

68. Olivera M, Delgado N, Cádiz F, Riquelme N, Montenegro I, Seeger M, Bravo G, Barros-Parada W, Pedreschi R, Besoain X (2021) Diffusible compounds produced by *Hanseniaspora osmophila* and *Gluconobacter cerinus* help to control the causal agents of gray rot and summer bunch rot of table grapes. Antibiotics 10: 664

69. Oukala N, Pastor-Fernández J, Sanmartín N, Aissat K, Pastor V (2021) Endophytic bacteria from the sahara desert protect tomato plants against *Botrytis cinerea* under different experimental conditions. Curr Microbiol 78: 2367–2379

70. Ownley BH, Duffy BK, Weller DM (2003) Identification and manipulation of soil properties to improve the biological control performance of phenazine-producing *Pseudomonas fluorescens*. Appl Environ Microbiol 69: 3333–3343

71. Prigigallo MI, Staropoli A, Vinale F, Bubici G (2023) Interactions between plant-beneficial microorganisms in a consortium: *Streptomyces microflavus* and *Trichoderma harzianum*. Microb. Biotechnol.

72. Quast C, Pruesse E, Yilmaz P, Gerken J, Schweer T, Yarza P, Peplies J, Glöckner FO (2012) The SILVA ribosomal RNA gene database project: improved data processing and web-based tools. Nucleic Acids Res 41: D590–D596

73. Ristaino JB, Anderson PK, Bebber DP, Brauman KA, Cunniffe NJ, Fedoroff NV, Finegold C, Garrett KA, Gilligan CA, Jones CM (2021) The persistent threat of emerging plant disease pandemics to global food security. Proc Natl Acad Sci 118: e2022239118

74. Rivers AR, Weber KC, Gardner TG, Liu S, Armstrong SD (2018) ITSxpress: software to rapidly trim internally transcribed spacer sequences with quality scores for marker gene analysis. F1000Research 7

75. Sandrini M, Nerva L, Sillo F, Balestrini R, Chitarra W, Zampieri E (2022) Abiotic stress and belowground microbiome: The potential of omics approaches. Int J Mol Sci 23: 1091

76. Shannon P, Markiel A, Ozier O, Baliga NS, Wang JT, Ramage D, Amin N, Schwikowski B, Ideker T (2003) Cytoscape: a software environment for integrated models of biomolecular interaction networks. Genome Res 13: 2498–2504

77. Sharp RE, LeNoble ME (2002) ABA, ethylene and the control of shoot and root growth under water stress. J Exp Bot 53: 33–37

78. Singh TB, Sahai V, Ali A, Prasad M, Yadav A, Shrivastav P, Goyal D, Dantu PK (2020) Screening and evaluation of PGPR strains having multiple PGP traits from hilly terrain. J Appl Biol Biotechnol 8: 38–44

79. Svenningsen NB, Watts-Williams SJ, Joner EJ, Battini F, Efthymiou A, Cruz-Paredes C, Nybroe O, Jakobsen I (2018) Suppression of the activity of arbuscular mycorrhizal fungi by the soil microbiota. ISME J 12: 1296–1307

80. Teale WD, Paponov IA, Palme K (2006) Auxin in action: signalling, transport and the control of plant growth and development. Nat Rev Mol Cell Biol 7: 847–859

81. Toju H, Peay KG, Yamamichi M, Narisawa K, Hiruma K, Naito K, Fukuda S, Ushio M, Nakaoka S, Onoda Y (2018) Core microbiomes for sustainable agroecosystems. Nat Plants 4: 247–257

82. Ton J, Flors V, Mauch-Mani B (2009) The multifaceted role of ABA in disease resistance. Trends Plant Sci 14: 310–317

83. Tufail MA, Ayyub M, Irfan M, Shakoor A, Chibani CM, Schmitz RA (2022) Endophytic bacteria perform better than endophytic fungi in improving plant growth under drought stress: A meta-comparison spanning 12 years (2010–2021). Physiol Plant 174: e13806

84. Van Huylenbroeck JM, van Laere IM, Piqueras A, Debergh PC, Bueno P (1998) Time course of catalase and superoxide dismutase during acclimatization and growth of micropropagated *Calathea* and *Spathiphyllum* plants. Plant Growth Regul 26: 7–14

85. Vandenkoornhuyse P, Quaiser A, Duhamel M, Le Van A, Dufresne A (2015) The importance of the microbiome of the plant holobiont. New Phytol 206: 1196–1206

86. Veach AM, Morris R, Yip DZ, Yang ZK, Engle NL, Cregger MA, Tschaplinski TJ, Schadt CW (2019) Rhizosphere microbiomes diverge among *Populus trichocarpa* plant-host genotypes and chemotypes, but it depends on soil origin. Microbiome 7: 1–15

87. Verhagen BW, Glazebrook J, Zhu T, Chang H-S, Van Loon L, Pieterse CM (2004) The transcriptome of rhizobacteria-induced systemic resistance in *Arabidopsis*. Mol Plant Microbe Interact 17: 895–908

88. Verhagen BW, Trotel-Aziz P, Couderchet M, Höfte M, Aziz A (2010) *Pseudomonas* spp.-induced systemic resistance to *Botrytis cinerea* is associated with induction and priming of defence responses in grapevine. J Exp Bot 61: 249–260

89. Viaene T, Langendries S, Beirinckx S, Maes M, Goormachtig S (2016) *Streptomyces* as a plant’s best friend? FEMS Microbiol. Ecol. 92

90. Wan M, Li G, Zhang J, Jiang D, Huang H-C (2008) Effect of volatile substances of *Streptomyces platensis* F-1 on control of plant fungal diseases. Biol Control 46: 552–559

91. Wei Y, Wu Y, Yan Y, Zou W, Xue J, Ma W, Wang W, Tian G, Wang L (2018) High-throughput sequencing of microbial community diversity in soil, grapes, leaves, grape juice and wine of grapevine from China. PLOS ONE 13: e0193097

92. Wiese R, Eiglsperger M, Kaufmann M (2004) yfiles—visualization and automatic layout of graphs. Graph Draw. Softw. Springer, pp 173–191

93. Yu K, Pieterse CM, Bakker PA, Berendsen RL (2019) Beneficial microbes going underground of root immunity. Plant Cell Environ 42: 2860–2870

94. Zou Y, Xue W, Luo G, Deng Z, Qin P, Guo R, Sun H, Xia Y, Liang S, Dai Y (2019) 1,520 reference genomes from cultivated human gut bacteria enable functional microbiome analyses. Nat Biotechnol 37: 179–185

